# Kinase-mediated RAS signaling via membraneless cytoplasmic protein granules

**DOI:** 10.1101/704312

**Authors:** Asmin Tulpule, Juan Guan, Dana S. Neel, Yone Phar Lin, Ann Heslin, Hannah R. Allegakoen, Shriya Perati, Alejandro D. Ramirez, Xiaoyu Shi, Bin Yang, Siyu Feng, Suraj Makhija, David Brown, Bo Huang, Trever G. Bivona

## Abstract

Receptor tyrosine kinase (RTK)-mediated activation of downstream effector pathways such as the RAS GTPase/MAP kinase (MAPK) signaling cascade is thought to occur exclusively from lipid membrane compartments in mammalian cells. Here, we uncover a membraneless, protein granule-based subcellular structure that can organize RTK/RAS/MAPK signaling in cancer. Chimeric (fusion) oncoproteins involving certain RTKs including ALK and RET undergo *de novo* higher-order assembly into membraneless cytoplasmic protein granules. These pathogenic biomolecular condensates locally concentrate the RAS activating complex GRB2/SOS1 and activate RAS in a lipid membrane-independent manner to initiate MAPK signaling. Formation of membraneless protein granules by RTK oncoproteins is both necessary and sufficient for RAS/MAPK signaling output in cells. Our findings reveal membraneless, higher-order cytoplasmic protein assembly as a distinct subcellular platform to activate RTKs and RAS GTPases and a general principle by which cells can organize oncogenic signaling.

**Highlights:**

- RTK oncoproteins can form *de novo* membraneless cytoplasmic protein granules
- RTK protein granules activate RAS in lipid membrane-independent manner
- Higher-order protein assembly is critical for oncogenic RAS/MAPK signaling
- Protein granules are a distinct subcellular platform for organizing RTK signaling

## Introduction

RTK/RAS/MAPK signaling is broadly important in regulating the proliferation and survival of normal human cells and is often hyper-activated through various mechanisms in human cancer (Sanchez-Vega et al., 2018). Native RTKs are integral membrane proteins and canonical RTK signaling is thought to occur exclusively from lipid-membrane subcellular compartments including the plasma membrane (PM) and intracellular organelles such as endosomes (Lemmon and Schlessinger, 2010). Moreover, RAS GTPase activation and downstream MAPK signaling is dependent on the lipid membrane association of RAS proteins (Cox et al., 2015; Willumsen et al., 1984). Evidence of local RTK and RAS protein clustering in PM lipid-microdomains (Delos Santos et al., 2015; Plowman et al., 2005; Prior et al., 2003) and recent reports that the PM resident T-cell receptor and associated proteins undergo phase separation in the presence of lipid bilayers highlights the importance of physical compartmentalization of signaling events (Huang et al., 2019; Su et al., 2016). Distinct from the PM and lipid membrane-bound organelles, biomolecular condensates are an emerging mechanism of subcellular compartmentalization through primarily protein-based membraneless structures such as P-bodies, nucleoli and stress granules (Alberti et al., 2019; Shin and Brangwynne, 2017). Though connections between aberrant transcription factor condensates and cancer have been proposed (Boulay et al., 2017; Koken et al., 1994), the functional role of biomolecular condensates in oncogenic signaling and cancer pathogenesis remains to be defined.

Prominent examples of oncogenic RTK/RAS/MAPK signaling in cancer include naturally occurring chromosomal rearrangements involving RTKs such as anaplastic lymphoma kinase (ALK) or rearranged during transfection (RET), which generate chimeric (fusion) oncoproteins that are validated therapeutic targets across multiple cancer subtypes (Childress et al., 2018; Kato et al., 2017). Virtually all oncogenic ALK and RET fusion proteins retain the intracellular domain, which includes the kinase, but lack the native transmembrane domain (Childress et al., 2018; Kato et al., 2017). The absence of a canonical lipid-membrane targeting domain as a shared structural feature of many oncogenic RTK fusion proteins presents a fundamental cell biological question (Nelson et al., 2017): how do these RTK fusion oncoproteins activate RAS signaling?

We previously discovered that the echinoderm microtubule-associated protein-like 4 (EML4)- ALK fusion oncoprotein that is present recurrently in lung cancer and other cancer subtypes is exquisitely dependent upon RAS GTPase activation and downstream RAF/MEK/ERK (MAPK pathway) signaling for its oncogenic output (Hrustanovic et al., 2015). We and other groups showed that EML4-ALK is not localized to the PM, but instead to intracellular, punctate cytoplasmic structures of unknown identity (Hrustanovic et al., 2015; Richards et al., 2015). This specific intracellular localization is essential for EML4-ALK to activate RAS and downstream MAPK signaling (Hrustanovic et al., 2015). Neither the biophysical or biochemical nature of these cytoplasmic structures nor the mechanism through which they promote oncogenic RAS signaling is clear. In this study, we uncover a previously unrecognized mode of RTK/RAS/MAPK signaling in mammalian cells that emanates from membraneless cytoplasmic protein granules.

## Results

### EML4-ALK forms *de novo* membraneless cytoplasmic protein granules

We set out to identify the cytoplasmic structure to which ALK fusion oncoproteins localize in cells. We focused our initial study on EML4-ALK variant 1, the most common oncogenic form in human cancers (Sabir et al., 2017). First, we confirmed that EML4-ALK localized to punctate structures in the cytoplasm, and not to the PM, by immunofluorescence (IF) in patient-derived cancer cells (H3122) that endogenously express this EML4-ALK variant (Hrustanovic et al., 2015) (Figure 1A). We validated the similar presence of EML4-ALK cytoplasmic puncta upon expression in a non-transformed human bronchial epithelial cell line (Beas2B), both by IF analysis in Beas2B cells expressing FLAG-tagged EML4-ALK (Figure S1A) and by live-cell imaging in Beas2B cells expressing fluorescent protein-tagged EML4-ALK (Figure S1B). These imaging results confirm localization of EML4-ALK at cytoplasmic puncta and indicate these subcellular structures are not the result of artificial expression, fixation, or fluorescent protein-mediated multimerization (Cranfill et al., 2016).

**Figure 1:**
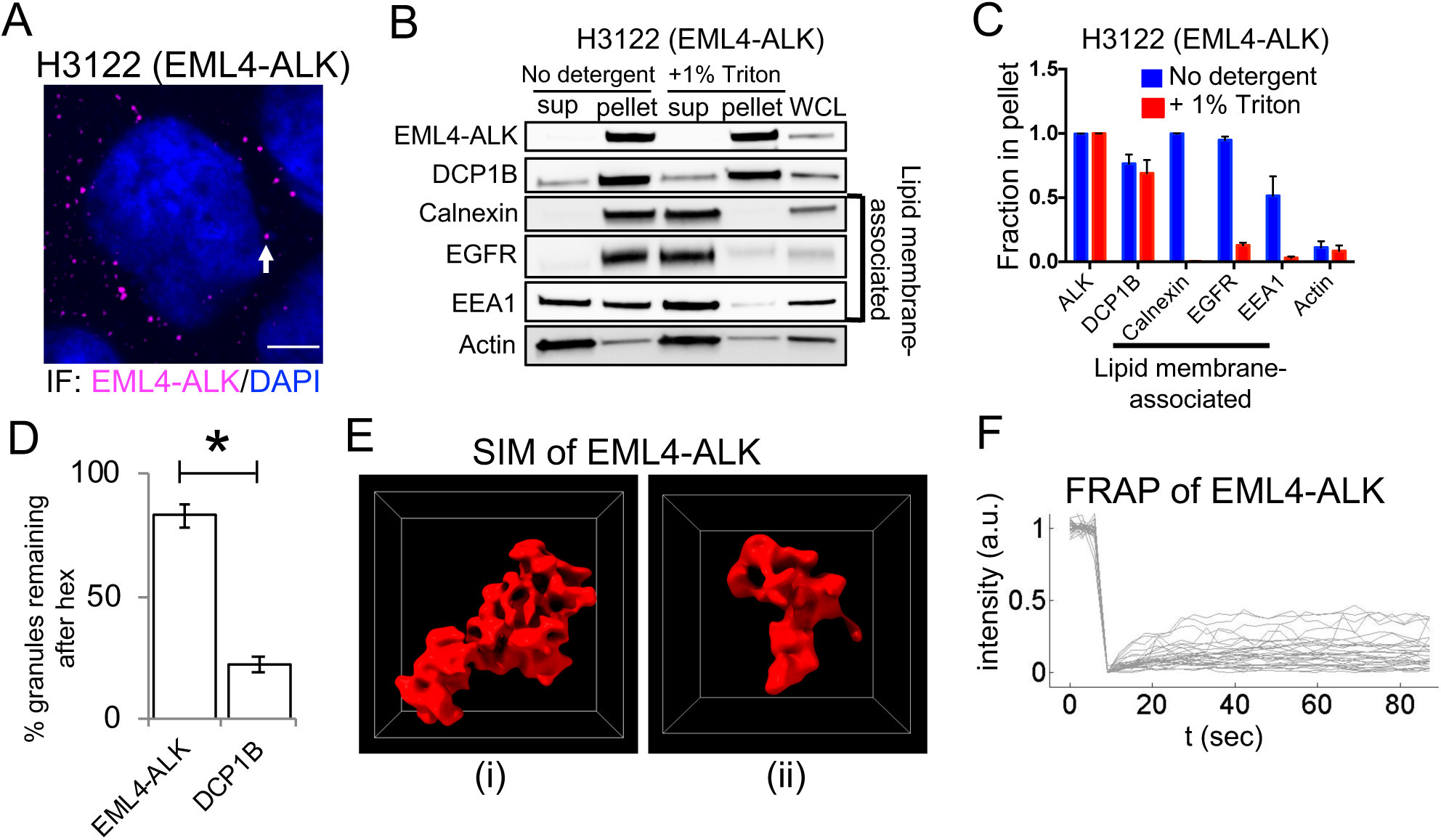
EML4-ALK forms *de novo* membraneless cytoplasmic protein granules. **(A)** Anti-ALK immunofluorescence (IF) to detect endogenous expression of EML4-ALK in a patient-derived cancer cell line (H3122). White arrow indicates a representative EML4-ALK cytoplasmic puncta. DAPI serves as a nuclear stain. Images are representative of at least 20 analyzed cells in 3 independent experiments. Scale bar = 5 µm. **(B, C)** Subcellular fractionation by ultracentrifugation +/– detergent (1% Triton X-100) in EML4-ALK expressing cancer cell line H3122, followed by Western blotting (B). EML4-ALK and DCP1B are statistically distinct (p < 0.05, one-way ANOVA) from the lipid membrane-associated proteins, which shift from the insoluble fraction (pellet) to the supernatant (sup) with detergent. Fraction in pellet (C) calculated as ratio of the insoluble fraction to total (insoluble plus supernatant fractions) as assessed by Western blotting, N = 3. **(D)** Quantification of EML4-ALK or DCP1B cytoplasmic granule persistence after 5 minutes of 5% hexanediol treatment. Error bars represent ± SEM, * denotes p value < 0.05 by unpaired *t*- test. **(E)** SIM images of 2 distinct YFP::EML4-ALK puncta in Beas2B cells. SIM box size: 2 µm × 2 µm × 2 µm. **(F)** FRAP analysis of YFP::EML4-ALK expressed in human epithelial cell line Beas2B. Each curve represents photobleaching and recovery of fluorescence intensity for an individual EML4- ALK puncta. N = 30 cells.

We next tested whether EML4-ALK cytoplasmic puncta correspond to an intracellular lipid membrane-containing structure, given the well-established role of lipid membranes in organizing RTK signaling and the requirement of lipid membranes for RAS GTPase activation (Delos Santos et al., 2015; Jackson et al., 1990; Willumsen et al., 1984). Live-cell imaging in Beas2B cells showed no significant colocalization of EML4-ALK cytoplasmic puncta with the PM or intracellular membranes as marked by a lipid membrane intercalating dye, or with a panel of established protein markers labeling canonical intracellular lipid-containing organelles (Rizzuto et al., 1995) (Figure S1C). To further evaluate whether EML4-ALK associates with lipid membranes, we performed subcellular fractionation in patient-derived cancer cell lines expressing endogenous EML4-ALK protein. EML4-ALK displayed a fractionation pattern unaffected by membrane-solubilizing detergents, which was distinct from the pattern of PM- spanning (epidermal growth factor receptor, EGFR) or internal membrane proteins (calnexin and early endosome antigen 1, EEA1), yet similar to that of a well-known cytoplasmic ribonucleoprotein granule constituent (the P-body protein de-capping mRNA 1B, DCP1B (Aizer et al., 2008)) (Figures 1B and 1C, and Figures S1D and S1E). These findings indicated that EML4-ALK may exist in a membraneless subcellular compartment within the cytoplasm. We confirmed by fluorescence microscopy that EML4-ALK puncta do not colocalize with the two known biomolecular condensates in the cytoplasm, P-bodies and stress granules (Figure S1C). EML4-ALK puncta are also not disrupted by RNase A, in contrast to ribonucleoprotein granules like the P-body (Figures S1F and S1G). These results suggest that EML4-ALK forms distinct protein-based, instead of RNA-protein-based, membraneless cytoplasmic granules.

We further investigated the biophysical nature of EML4-ALK cytoplasmic granules using a suite of established cellular assays to characterize biomolecular condensates (Molliex et al., 2015; Patel et al., 2015). During live-cell imaging, no fission or fusion of EML4-ALK granules was observed in spite of occasional granule collisions (Video S1), which is in contrast to the expected behaviors for liquid-like granules (Alberti et al., 2019). Unlike DCP1B-labelled P-bodies, we found that EML4-ALK granules mostly persist after hexanediol treatment, which disrupts many liquid-like condensates (Kroschwald et al., 2015) (Figure 1D, and Figure S1H). Super-resolution imaging by Structured Illumination Microscopy (SIM) revealed that the EML4-ALK granules exhibit porous and curvilinear features that are distinct from the more smooth and spherical appearance of liquid-like granules (Patel et al., 2015; Shin and Brangwynne, 2017) (Figure 1E). Moreover, fluorescence recovery after photo-bleaching (FRAP) showed an overall low fraction of exchange of EML4-ALK between the granules and the surrounding cytosol (Figure 1F). This recovery fraction was heterogeneous amongst granules, varying from 40% to negligible recovery at 1 minute, possibly reflecting an ongoing aging process as maturing granules adopt increasingly solid-like states (Molliex et al., 2015). Taken together, the data indicate EML4-ALK forms *de novo* membraneless cytoplasmic protein granules that are distinct from well-known cytoplasmic biomolecular condensates and demonstrate biophysical properties that are more solid than liquid-like.

### EML4-ALK membraneless cytoplasmic protein granules recruit the RAS-activating complex GRB2/SOS1/GAB1 *in situ*

To uncover the connection between EML4-ALK membraneless cytoplasmic granules and RAS activation, we created a library of gene-edited Beas2B cell lines by introducing a split mNeonGreen2_1-10/11_ tag (mNG2) at the endogenous locus of canonical adaptor and effector proteins in the RTK/RAS/MAPK signaling pathway, including GRB2, GAB1, SOS1, and RAS GTPases (H/N/K isoforms) (Feng et al., 2017). This suite of isogenic cell lines avoids potential biases that can arise when overexpressing labeled proteins or fixing and permeabilizing cells for immunofluorescence. In this set of cell lines, we found that expression of EML4-ALK specifically re-localized key upstream RAS pathway proteins, including GRB2, GAB1, and SOS1, from a mainly cytosolic pattern to the discrete EML4-ALK granules, and not to the PM (Figures 2A and 2B). This is distinct from the pattern of PM re-localization seen in the control case of expressing an oncogenic form of the transmembrane RTK EGFR (Figure S2A). Treatment with the ALK kinase inhibitor crizotinib for 24 hours substantially reduced the recruitment of these adaptor proteins without affecting cell viability, indicating that this process requires ALK kinase activation (Figure 2C). We orthogonally confirmed recruitment of the key adaptor, GRB2, to endogenous EML4-ALK cytoplasmic protein granules detected by IF in patient-derived cancer cells (H3122) (Figure 2D), as well as through dual expression of EML4- ALK and GRB2 in Beas2B cells (Figure S2B). Additionally, we observed a low and heterogeneous FRAP recovery behavior for GRB2 at the EML4-ALK protein granules, similar to that of EML4-ALK itself (Figure S2C).

**Figure 2:**
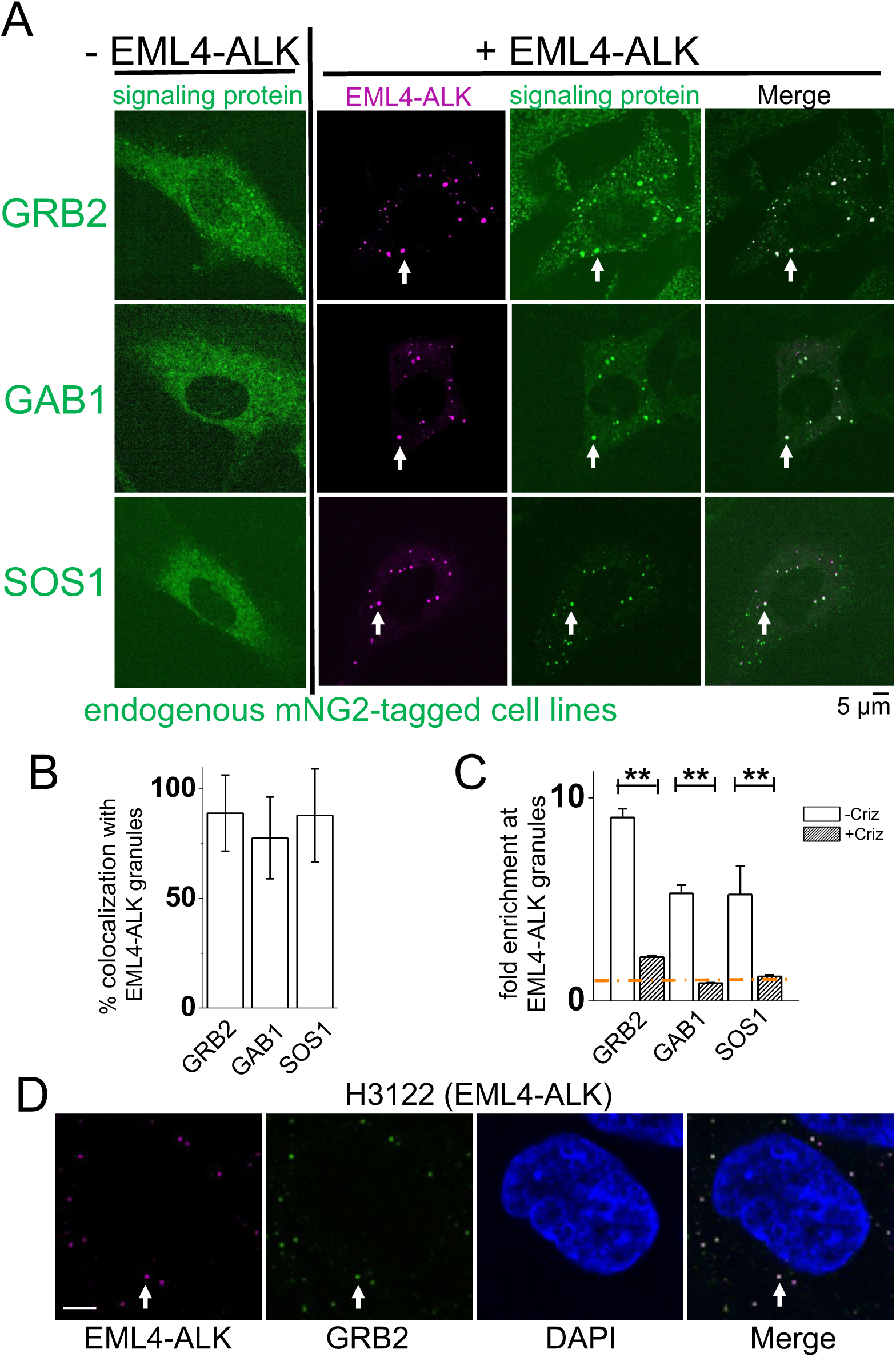
EML4-ALK membraneless cytoplasmic protein granules recruit RAS-activating complex GRB2/SOS1/GAB1 *in situ*. **(A)** Live-cell confocal imaging in Beas2B cells with endogenous mNG2-tagging of GRB2, GAB1, and SOS1 in the presence or absence of mTagBFP2::EML4-ALK. White arrows indicate a representative EML4-ALK cytoplasmic protein granule with local enrichment of respective signaling proteins (multiple non-highlighted granules also show colocalization between EML4- ALK and signaling proteins). **(B)** Quantification of colocalization between EML4-ALK granules and relevant signaling proteins. At least 100 total cells were scored in each condition over 3 independent experiments. Error bars represent ± SEM. **(C)** Fold-enrichment of signaling proteins at EML4-ALK granules +/– 24 hour treatment with 1 µM crizotinib (criz). Error bars represent ± SEM, ** p < 0.01, paired *t*-test. Orange dotted line at Y = 1 denotes level of zero enrichment. **(D)** Immunofluorescence to detect endogenous expression of EML4-ALK in a patient-derived cancer cell line (H3122) with endogenous mNG2-tagging of GRB2. White arrows indicate a representative EML4-ALK cytoplasmic protein granule with local enrichment of GRB2 (multiple non-highlighted granules also show colocalization between EML4-ALK and GRB2). Scale bar = 5 µM.

### Cytoplasmic EML4-ALK protein granules locally activate RAS

Our imaging and biochemical data prompted the unanticipated hypothesis that RTK-mediated RAS GTPase activation may occur via a subcellular structure lacking lipid membranes (i.e. EML4-ALK membraneless cytoplasmic protein granules), potentially through a cytosolic pool of RAS that is known to exist but with unclear functional significance (Goodwin et al., 2005; Zhou et al., 2016). We first confirmed RAS protein expression in the cytosol, in addition to lipid membrane subcellular compartments (Figure S3A). Next, we directly tested whether cytosolic RAS can become activated in a lipid membrane-independent manner by EML4-ALK cytoplasmic protein granules. We utilized established mutant forms of RAS (KRAS-C185S, H/NRAS-C186S) that abrogate lipid membrane targeting and are retained exclusively in the cytosol (Jackson et al., 1990). While the expression of either EML4-ALK or the PM-localized oncogenic EGFR increased RAS-GTP levels (Figure 3A, and Figures S3B and S3D), only EML4-ALK increased RAS-GTP levels of cytosolic RAS mutants (Figure 3B, and Figures S3C and S3E). Furthermore, inhibition of EML4-ALK with crizotinib in H3122 patient-derived cancer cells suppressed not only wild-type RAS-GTP levels, but also the levels of GTP-bound, cytosolic KRAS-C185S (Figures 3C and 3D). Control experiments treating a distinct patient-derived cancer cell line HCC827 expressing endogenous oncogenic EGFR (PM-localized) with an established EGFR inhibitor (Tsao et al., 2005) confirmed suppression of wild-type RAS-GTP levels but showed no effect on KRAS-C185S RAS-GTP levels (Figures S3F and S3G). These findings demonstrate the specificity of cytosolic RAS activation by oncogenic EML4-ALK.

**Figure 3:**
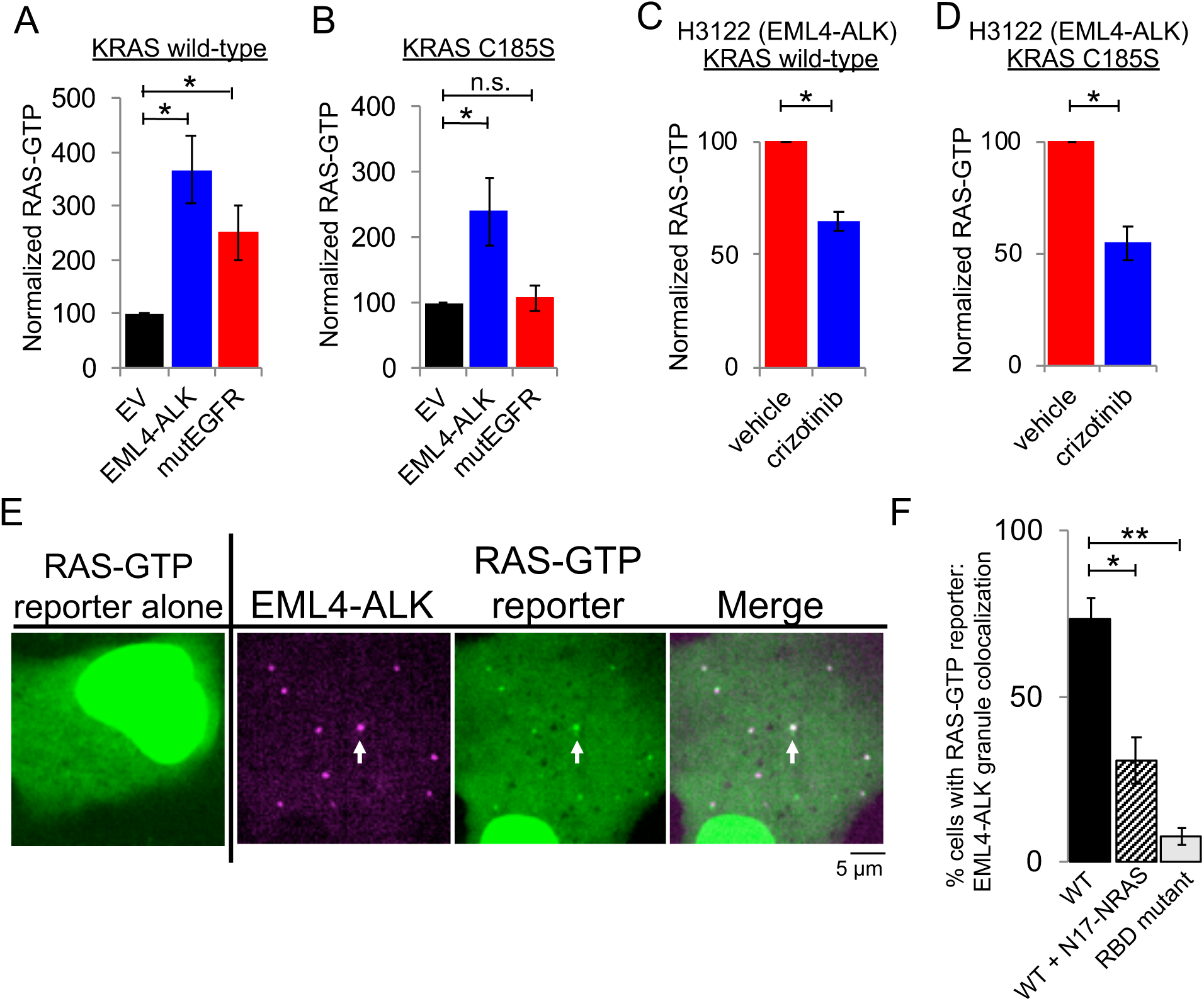
Cytoplasmic EML4-ALK protein granules locally activate RAS. **(A, B)** Stable expression of KRAS wild-type (A) or C185S cytosolic mutant (B) in 293T cells, followed by transfection of empty vector (EV), EML4-ALK, or oncogenic EGFR. RAS-GTP levels normalized to relevant total RAS species (KRAS wild-type or C185S) and then standardized against EV, N = 3. **(C, D)** EML4-ALK expressing H3122 cancer cell line with stable expression of KRAS wild-type (C) or cytosolic KRAS C185S mutant (D) +/– two hours of 100 nM crizotinib. RAS-GTP levels normalized to relevant total RAS species (KRAS wild-type or C185S) and then standardized against DMSO treated H3122 cells (vehicle), N = 3. **(E)** Live-cell confocal imaging of RAS-GTP reporter (tandem GFP-RBD) expressed in human epithelial cell line Beas2B +/- mTagBFP2::EML4-ALK. White arrows indicate a representative EML4-ALK cytoplasmic protein granule with local enrichment of RAS-GTP (multiple non-highlighted granules also show colocalization between EML4-ALK and RAS-GTP reporter). **(F)** Quantification of cells with colocalization between RAS-GTP reporter and EML4-ALK granules. WT denotes unmodified tandem GFP-RBD reporter, RBD mutant denotes mutant GFP-RBD reporter (R59A/N64D) with diminished RAS-GTP binding. N = 3 with at least 30 cells per replicate. **For all panels**, error bars represent ± SEM, * denotes p < 0.05, ** p < 0.01, n.s. denotes non-significant comparison, one-way ANOVA (A, B, F) or paired *t*-test (C, D).

Lastly, to determine whether EML4-ALK cytoplasmic granules display evidence of local RAS activation (i.e. RAS-GTP), we used an established tandem GFP-RBD (RAS-binding domain) live-cell reporter given its high affinity binding to RAS-GTP and sensitivity for detection of endogenous RAS activation (Biskup and Rubio, 2014). When expressed alone, the RAS-GTP reporter displayed homogenous localization in the cytosol and enrichment in the nucleoplasm, as previously described (Rubio et al., 2010) (Figure 3E). As a positive control, expression of oncogenic KRAS in Beas2B cells led to re-localization of the RAS-GTP reporter to the PM (Figure S3H). In EML4-ALK expressing cells, we observed robust enrichment of the RAS-GTP reporter at EML4-ALK cytoplasmic protein granules and not at the PM (Figures 3E and 3F). Co-expression of a dominant negative RAS (RASN17) (Sigal et al., 1986) that interferes with RAS activation (GTP-loading) decreased colocalization of the RAS-GTP reporter at EML4-ALK granules, as did introduction of mutations into the RBD component of the GFP-RBD reporter (RBD R59A/N64D) that decrease affinity for RAS-GTP (Biskup and Rubio, 2014) (Figure 3F). The collective findings show that local RAS activation and accumulation of RAS-GTP occurs at membraneless EML4-ALK cytoplasmic protein granules.

### Protein granule formation by EML4-ALK is critical for RAS/MAPK signaling

We next tested whether downstream MAPK signaling output is dependent on EML4-ALK cytoplasmic protein granules by investigating the molecular determinants of *de novo* granule formation. The EML4 portion of the chimeric EML4-ALK oncoprotein contains an N-terminal trimerization domain (TD) and a truncated tandem atypical WD-propeller in EML4 protein (TAPE) domain (Sabir et al., 2017) (Figure 4A). Deletion of the TD or the hydrophobic EML protein (HELP) motif in the propeller domain disrupted protein granule formation, resulting instead in a diffuse cytosolic distribution of EML4-ALK labeled by either fluorescent protein or FLAG-tag (Figures 4B and 4C, and Figure S4A). ΔTD or ΔHELP mutants of EML4-ALK demonstrated loss of ALK trans-phosphorylation and GRB2 interaction (Figure 4D, and Figure S4B) and impaired RAS/MAPK activation (Figures 4E and 4F, and Figure S4B). These data implicate *de novo* protein granule formation that is mediated by the EML4 portion of the fusion protein as critical for productive RAS/MAPK signaling. An established kinase-deficient mutant (K589M) form of EML4-ALK demonstrated diminished RAS/MAPK signaling and also disrupted protein granule formation (Figures 4B-4F, and Figures S4A and S4B), an effect that may be due to phosphorylation events regulating EML4-ALK protein granule assembly, as has been observed with other biomolecular condensates (Monahan et al., 2017; Rai et al., 2018).

**Figure 4:**
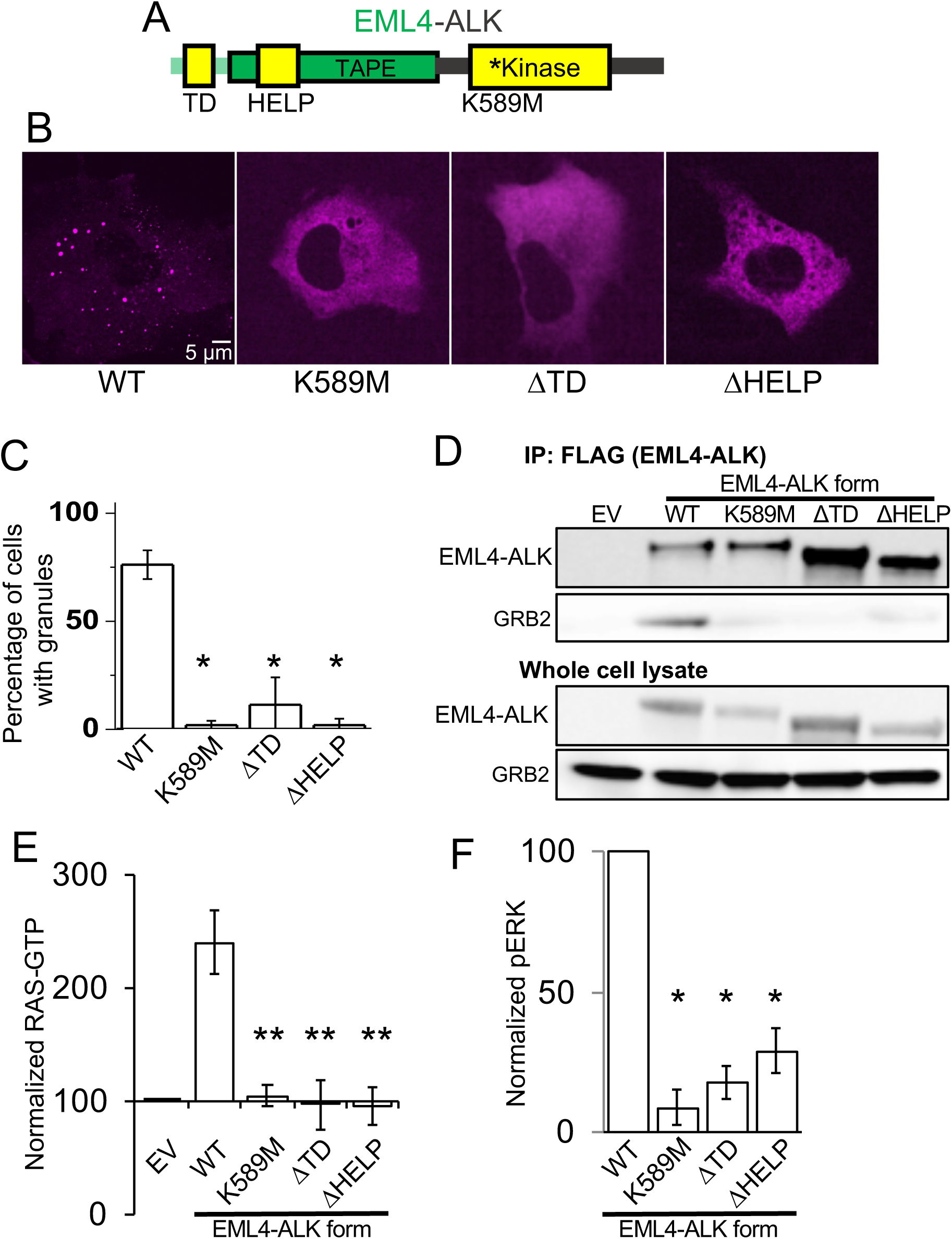
Protein granule formation by EML4-ALK is critical for RAS/MAPK signaling. **(A)** Structure schematic of the EML4-ALK fusion protein with highlighted trimerization domain (TD), hydrophobic EML protein (HELP) motif within the tandem atypical WD-propeller in EML4 protein (TAPE) domain, and ALK kinase domain. **(B)** Live-cell confocal imaging of mTagBFP2::EML4-ALK (denoted as WT for wild-type EML4-ALK) or kinase-deficient (K589M), ΔTD or ΔHELP mutants in human epithelial cell line Beas2B. **(C)** Quantification of percentage of cells with granules. 75 cells were scored for each condition over 3 independent replicates. **(D)** Anti-FLAG co-immunoprecipitation of FLAG-tagged wild-type (WT) EML4-ALK or respective mutants expressed in 293T cells, followed by Western blotting to assess GRB2 binding. EV denotes empty vector control, images representative of at least 3 independent experiments. **(E, F)** Quantification of endogenous RAS-GTP levels (E) and ERK activation (F) by Western blotting upon expression of EML4-ALK or respective mutants in 293T cells. RAS-GTP levels were normalized to total RAS protein levels and then displayed relative to EV sample which was set to 100, N = 4. pERK levels were normalized to total ERK and then displayed relative to wild- type EML4-ALK sample which was set to 100, N = 4. **For all panels**, error bars represent ± SEM, * denotes p < 0.05, ** denotes p < 0.01, one-way ANOVA.

### Higher-order clustering of an RTK in membraneless cytoplasmic protein granules is sufficient to activate RAS/MAPK signaling

Biomolecular condensates, such as EML4-ALK protein granules, are typically micron-sized, membraneless, higher-order protein assemblies (Banani et al., 2017). Using structural mutants of EML4-ALK, we demonstrated that disruption of protein granule formation impaired RAS/MAPK pathway activation. We therefore hypothesized that higher-order clustering of an RTK in membraneless cytoplasmic protein granules is sufficient to organize activation of RAS/MAPK signaling. To directly test this hypothesis, we utilized the HOtag method developed recently to enable forced protein granule formation through multivalent interactions that allow for higher-order protein assembly (Zhang et al., 2018) (Figure S5A). HOtag-induced cytoplasmic granule formation of either the ΔTD or ΔHELP mutants of EML4-ALK locally recruited GRB2 (Figures 5A and 5B), increased RAS-GTP levels (Figure S5B) and restored RAS/MAPK signaling (Figures 5C and 5D). As an important negative control, HOtag-forced clustering of the kinase-deficient EML4-ALK did not promote GRB2 recruitment or RAS/MAPK signaling (Figures 5A-5D, and Figure S5B). The findings highlight the dual importance of cytoplasmic protein granule formation and intact kinase activity for productive signaling. We also directly tested the role of protein granule formation on cytosolic RAS activation. Compared to wild-type EML4-ALK, the ΔTD mutant that is distributed diffusely in the cytosol demonstrated substantially reduced levels of activated (GTP-bound) cytosolic KRAS-C185S, which could be restored through HOtag-forced clustering (Figure S5C). Collectively, our data show that membraneless EML4-ALK cytoplasmic protein granules can spatially concentrate, organize, and initiate RAS/MAPK pathway signaling events.

**Figure 5:**
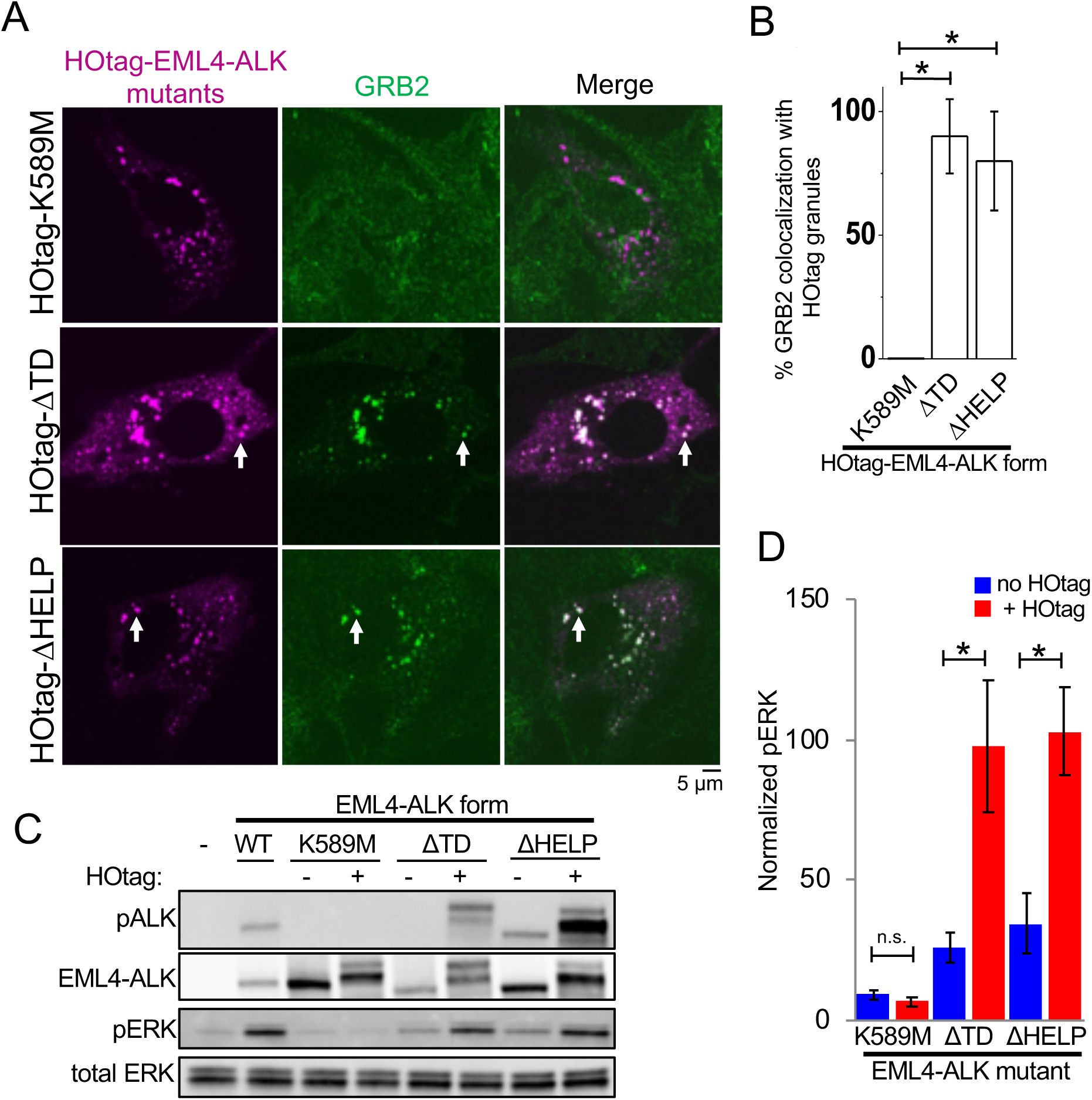
Forced clustering of EML4-ALK cytosolic mutants drives RAS/MAPK signaling. **(A)** Live-cell confocal imaging of HOtag-mTagBFP2::EML4-ALK ΔTD, ΔHELP, and K589M mutants in Beas2B cells with endogenous mNG2-tagging of GRB2. White arrows indicate representative HOtag-EML4-ALK ΔTD or ΔHELP protein granules with local enrichment of GRB2 (multiple non-highlighted granules also show colocalization between HOtag-EML4-ALK mutants and GRB2). **(B)** Quantification of percent colocalization between HOtag protein granules of EML4-ALK mutants and GRB2. N = 130 total cells for each condition over 3 independent experiments. **(C, D)** Western blotting upon expression of wild-type EML4-ALK or respective mutants +/– HOtag in 293T cells. For quantification, pERK levels were normalized to total ERK and then displayed relative to wild-type EML4-ALK sample which was set to 100, N = 5. **For all panels**, error bars represent ± SEM, * denotes p < 0.05, n.s. denotes non-significant comparison, one-way ANOVA (B) or paired *t-*test (D).

### Cytoplasmic granule formation is a general mechanism for oncogenic RTK-mediated RAS/MAPK pathway activation

We tested the generality of this model. First, multiple variants of EML4-ALK have been described in cancer patients (Sabir et al., 2017), all comprising the intracellular domain of ALK (but not its transmembrane domain) fused to N-terminal fragments of EML4 of varying lengths (Figure 6A). We demonstrated that another recurrent form of oncogenic EML4-ALK (variant 3), which contains a further truncation of the TAPE domain but retains the TD (Sabir et al., 2017), also formed cytoplasmic granules that locally recruited GRB2 and increased RAS/MAPK signaling (Figures 6B-6D, and Figures S6A-S6C). In contrast, EML4-ALK variant 5, which lacks the entire TAPE domain of EML4, did not form protein granules and demonstrated substantially less RAS/MAPK signaling compared to the protein granule-forming EML4-ALK variants 1 and 3 (Figures 6B-6D, and Figure S6D). HOtag-forced clustering of EML4-ALK variant 5 augmented RAS/MAPK signaling (Figures S6E and S6F). Consistent with the presence of a TD in all EML4-ALK variants, the granule-forming EML4-ALK variants (1 and 3) and the non-granule-forming variant 5 were each capable of self-association in co-immunoprecipitation experiments (Figure S6G). These results reinforce the role of higher-order protein assembly in promoting oncogenic signaling by these fusion RTKs. The findings suggest a functional difference in terms of RAS/MAPK signaling output between higher-order protein granule formation (EML4-ALK variants 1 and 3) and lower-order self-association through oligomerization (EML4-ALK variant 5).

**Figure 6:**
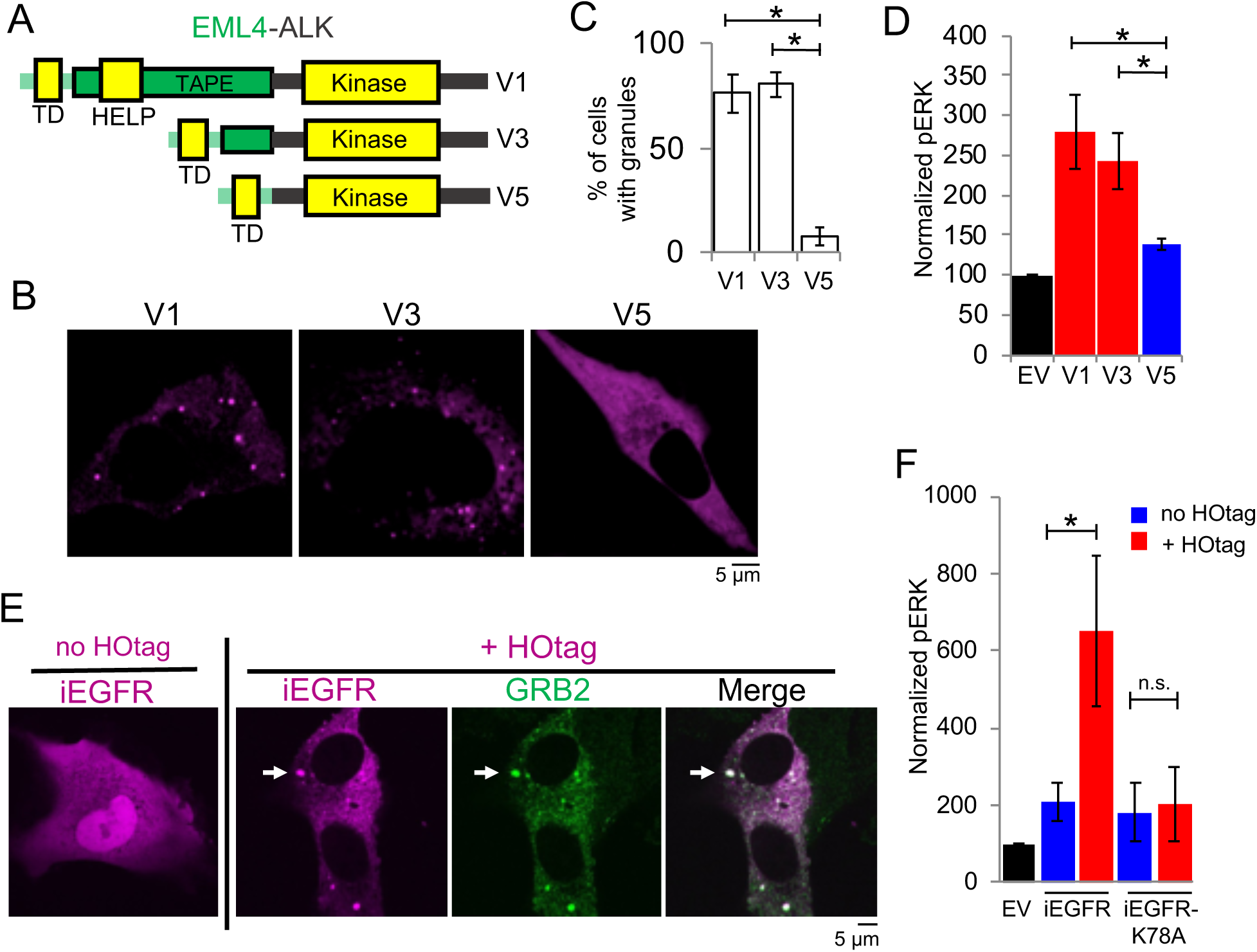
Higher-order protein assembly of RTK fusion oncoproteins drives oncogenic RAS/MAPK signaling. **(A)** Structure schematic of EML4-ALK fusion protein variants 1, 3, and 5 with highlighted trimerization domain (TD), HELP motif, and TAPE domain. **(B)** Live-cell confocal imaging of YFP::EML4-ALK variants 1, 3, and 5 in Beas2B cells. **(C)** Quantification of percentage of cells with granules. At least 100 cells were scored over 3 independent replicates. **(D)** Quantification of Western blotting results upon expression of EML4-ALK variants or empty vector (EV) in 293T cells. For quantification, pERK levels were normalized to total ERK and then displayed relative to EV sample which was set to 100, N = 3. **(E)** Live-cell confocal imaging of mTagBFP2::iEGFR +/– forced clustering (HOtag) in Beas2B cells with endogenous mNG2-tagging of GRB2. White arrows indicate a representative HOtag- iEGFR protein granule with local enrichment of GRB2 (multiple non-highlighted granules also show colocalization between HOtag-iEGFR and GRB2). **(F)** Quantification of Western blotting results upon expression of EV, iEGFR or iEGFR kinase-deficient mutant (iEGFR-K78A) +/– HOtag in 293T cells. For quantification, pERK levels were normalized to total ERK and then displayed relative to EV sample which was set to 100, N = 6. **For all panels**, images are representative of at least 25 analyzed cells in 3 independent experiments. Error bars represent ± SEM, * denotes p < 0.05, n.s. denotes non-significant comparison, one-way ANOVA (C, D) or paired *t-*test (F).

Next, as a proof-of-principle for the functional importance of higher-order protein assembly, we engineered an intracellular EGFR (iEGFR) protein lacking the native extracellular and transmembrane domains. This iEGFR is similar to naturally-occurring truncated forms of this RTK and others (Liao and Carpenter, 2012; Ni et al., 2001) and is distributed diffusely in the cytoplasm and nucleus when expressed alone (Figure 6E). HOtag-forced clustering of iEGFR recruited GRB2 and increased RAS/MAPK signaling in a kinase-dependent manner, analogous to oncogenic ALK (Figures 6E and 6F).

Furthermore, we studied another oncogenic RTK, RET, that also undergoes multiple distinct and recurrent gene rearrangements in human cancer, leading to the elimination of the extracellular and transmembrane domains from the various fusion oncoproteins (Kato et al., 2017). The fusion oncoprotein CCDC6-RET formed *de novo* cytoplasmic protein granules which did not demonstrate PM localization or colocalize with intracellular lipid-containing organelles or a lipid-intercalating dye (Figures 7A and 7B, and Figure S7A). CCDC6-RET cytoplasmic protein granules recruited GRB2 (Figure 7B) and locally enriched RAS-GTP as measured by the tandem GFP-RBD reporter (Figure 7C), resulting in increased RAS activation and downstream MAPK signaling (Figures 7D and 7E, and Figure S7B). Structure-function studies showed that a CCDC6-RET mutant lacking the coiled-coil domain in the CCDC6 component abrogated granule formation (Figure S7C) and reduced RAS/MAPK activation (Figures 7D and 7E, and Figure S7B). A kinase-deficient (K147M) mutant form of CCDC6-RET still formed cytoplasmic protein granules, yet was unable to recruit GRB2 (Figure S7D) or activate RAS/MAPK signaling (Figures 7D and 7E, and Figure S7B). These results reinforce the dual importance of cytoplasmic protein granules and kinase activity in driving oncogenic RTK/RAS/MAPK signaling. The data also reveal differences between RTK fusion oncoprotein subtypes (i.e. EML4-ALK and CCDC6- RET) in the dependence on kinase activity for granule formation.

**Figure 7:**
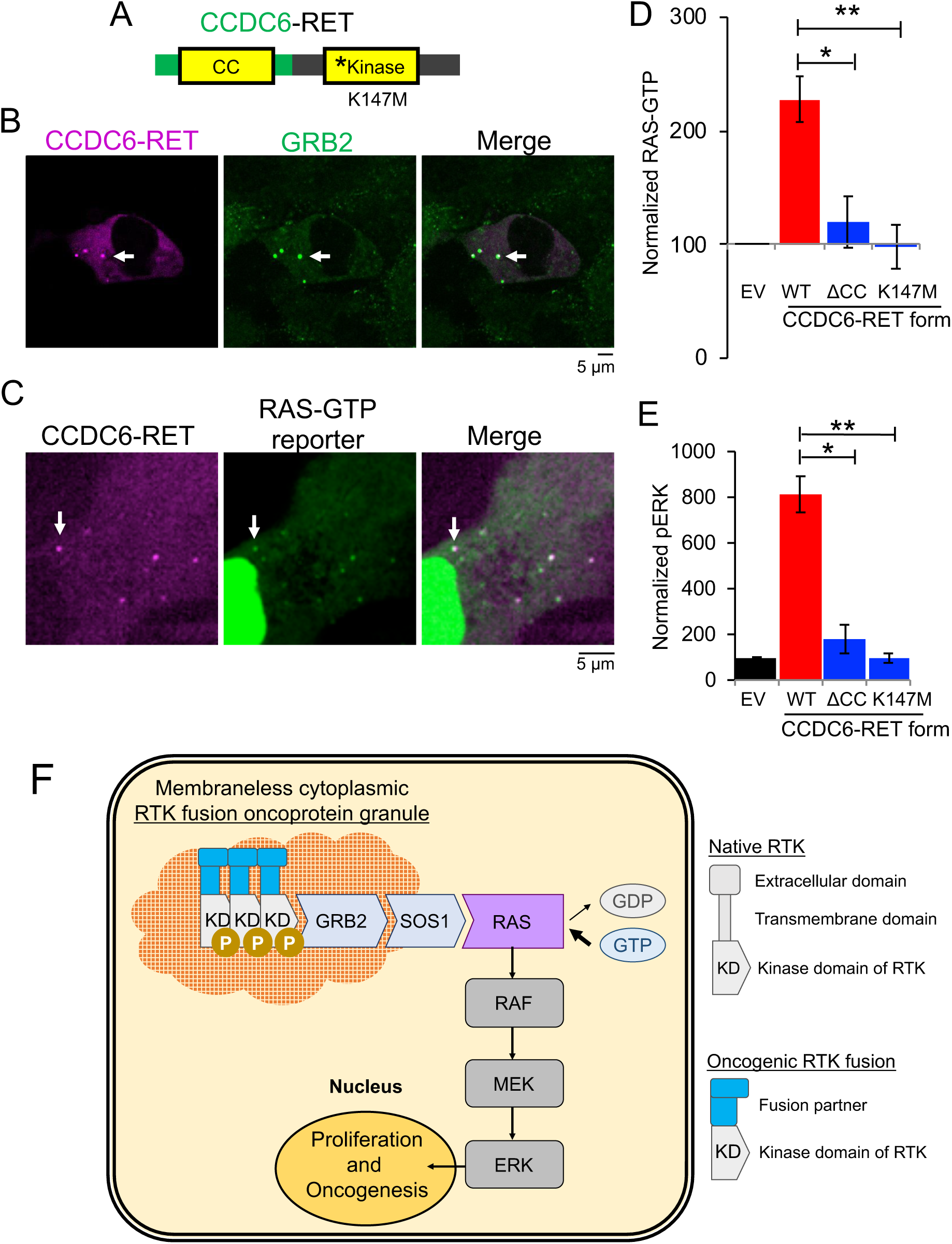
Cytoplasmic granule formation is a general mechanism for RTK-mediated RAS/MAPK pathway activation in cancer. **(A)** Structure schematic of the CCDC6-RET fusion protein with CCDC6 coiled-coiled domain (CC) and RET kinase domain. **(B)** Live-cell confocal imaging of mTagBFP2::CCDC6-RET in Beas2B cells with endogenous mNG2-tagging of GRB2. White arrows indicate a representative CCDC6-RET cytoplasmic protein granule with local enrichment of GRB2 (multiple non-highlighted granules also show colocalization between CCDC6-RET and GRB2). **(C)** Live-cell imaging of human epithelial cell line Beas2B expressing mTagBFP2::CCDC6-RET and tandem GFP-RBD (RAS-GTP reporter). White arrows indicate a representative CCDC6- RET cytoplasmic protein granule with local enrichment of RAS-GTP reporter (multiple non-highlighted granules also show colocalization between CCDC6-RET and RAS-GTP reporter). **(D, E)** Quantification of endogenous RAS-GTP levels (D) and ERK phosphorylation (E) by Western blotting in 293T cells expressing an empty vector (EV), CCDC6-RET wild-type (WT) or CCDC6-RET mutants (ΔCC and kinase-deficient mutant K147M). RAS-GTP levels were normalized to total RAS protein levels, and then displayed relative to EV sample which was set to 100. pERK levels were normalized to total ERK and then displayed relative to EV sample which was set to 100. N = 4. **(F)** Model for membraneless cytoplasmic protein granule-based oncogenic RTK/RAS/MAPK signaling. **For all panels**, images are representative of at least 25 analyzed cells in 3 independent experiments. Error bars represent ± SEM, * denotes p < 0.05, ** p < 0.01, n.s. denotes non-significant comparison, one-way ANOVA.

## Discussion

Collectively, our findings reveal a new mechanism for RTK activation and RAS signaling in cancer. We demonstrate that certain RTK fusion oncoproteins assemble *de novo* their own subcellular compartment, membraneless cytoplasmic protein granules, which coordinate local RAS activation in a lipid membrane-independent manner to drive oncogenic signaling (Figure 7F).

Physiologic RTKs, as well as oncogenic RTKs with kinase-activating missense or small insertion/deletion mutations, are integral membrane proteins that localize to and organize signaling events at lipid membrane subcellular compartments, including the PM and certain intracellular organelles (Lemmon and Schlessinger, 2010). In contrast, RTKs that undergo chromosomal rearrangements in cancer often lose their lipid membrane-targeting sequences (i.e. transmembrane domains) from the RTK fusion oncoprotein (Du and Lovly, 2018). The non-kinase fusion partner frequently contains multimerization domains which are important for self-association and oncogenic signaling (e.g. TD within EML4 in EML4-ALK) (Du and Lovly, 2018; McWhirter et al., 1993; Soda et al., 2007). However, it was not known whether RTK fusion oncoproteins form higher-order protein assemblies or whether multimerization alone was sufficient to drive oncogenic signaling. Here we report initial examples of RTK fusion oncoproteins forming biomolecular condensates that are critical for oncogenic RTK/RAS signaling. In the case of EML4-ALK, by testing multiple structural variants we determined that the presence of a multimerization domain alone is not sufficient for protein granule formation (non-granule-forming variant 5 contains a TD). This suggests that additional multivalent interactions are required for higher-order protein assembly. Moreover, variants of EML4-ALK that are capable of higher-order protein granule assembly displayed a significant functional difference (increased signaling output) compared to forms of EML4-ALK that were only competent for lower-order self-association.

We propose that *de novo* assembly of membraneless cytoplasmic protein granules may be a general mechanism for activating RTK fusion oncoprotein signaling in cancer. This mechanism is distinct from other known strategies of RTK activation including promoter driven overexpression of the oncoprotein or dimerization/oligomerization mediated by domains within the non-kinase fusion partner (Du and Lovly, 2018; Medves and Demoulin, 2012). We anticipate that general principles governing whether individual RTK fusion oncoproteins form membraneless cytoplasmic protein granules or other biomolecular condensates will emerge from further studies. Structural and biophysical features contributed by the non-kinase fusion partner including the number of multivalent interaction domains and the effect on intrinsic solubility may be important determinants of condensate formation by RTK fusion oncoproteins, as described in other protein granule systems (Banani et al., 2017). Post-translational modifications, such as phosphorylation, may regulate these multivalent interactions and enable the recruitment of adaptor proteins that affect condensate formation and stability. Interestingly, we found that *de novo* formation of ALK, but not RET, fusion oncoprotein assemblies require the kinase activity of the protein. This suggests that there may also be biochemical and structural differences amongst RTK fusion oncoprotein granules that will be important to determine.

Downstream of RTKs, RAS activation is a pathogenic hallmark of many cancers driven by chimeric RTK oncoproteins (Rotow and Bivona, 2017). The current paradigm holds that RAS proteins (e.g. H/N/K-RAS) are activated and signal to effector proteins such as the RAF/MEK/ERK kinases exclusively from lipid membrane compartments in mammalian cells (Cox et al., 2015; Willumsen et al., 1984). How naturally occurring chimeric RTK oncoproteins that lack lipid membrane targeting domains spatially coordinated RAS activation remained unclear. Our findings provide initial examples of RAS activation and productive downstream signaling from a membraneless subcellular compartment in mammalian cells. These results offer an alternative solution by which cells can organize oncogenic RTK/RAS/MAPK signaling that is distinct from canonical lipid membrane platforms such as the PM.

Subcellular compartments allow for the selective recruitment and interaction of signaling proteins, affording cells an additional layer of spatial control over the biochemical reactions driving signaling outputs (Chiu et al., 2002; Kim et al., 2007; Stoeger et al., 2016). For example, the altered dynamics of RTK/RAS signaling complexes at internal lipid membrane organelles may enable cells to modulate the intensity and duration of MAPK signaling (Rocks et al., 2005). Here we describe an alternative strategy to spatially concentrate signaling proteins and hyperactivate RTK/RAS/MAPK signaling through the formation of a pathogenic biomolecular condensate. Given the central role of RTK/RAS/MAPK signaling in both normal human development and numerous pathologies, a model of spatial regulation across different subcellular compartments including cytoplasmic protein granules could further explain how RAS GTPases as binary molecular switches can produce such pleiotropic and diverse signaling outputs in mammalian cells and disease pathologies (Fehrenbacher et al., 2009). The potential existence and role of membraneless higher-order protein assemblies in non-transformed cells as an additional subcellular compartment regulating physiologic RTK/RAS signaling is an area for future investigation.

The full complement of signaling proteins at membraneless cytoplasmic protein granules remains to be elucidated. RTKs can utilize an array of adaptor and effector proteins to regulate outputs from multiple signaling pathways (Lemmon and Schlessinger, 2010). The signaling protein architecture of RTK membraneless cytoplasmic granules may differ in important ways from lipid membrane-based RTK signaling complexes. It is possible that RTK signaling from protein granules preferentially engages with or accelerates specific signaling effectors such as the RAS/MAPK pathway. One hypothesis for this selectivity is that the biophysical properties of cytoplasmic protein granules (e.g. the solid-like state of EML4-ALK granules with minimal exchange of the adaptor protein GRB2) may sequester specific signaling proteins and alter the equilibrium between cytoplasmic granule and lipid membrane-based signaling protein pools. By increasing the local concentration of the RTK itself, as well as RAS adaptor proteins, cytoplasmic protein granules may shift the balance of RAS GTP/GDP exchange towards RAS activation and MAPK signaling in cancer cells. Alternatively, certain proteins regulating RTK/RAS signaling such as specific RAS guanine exchange factor (GEF) complex proteins and/or RAS GTPase activating proteins could be enriched or dis-enriched at the membraneless cytoplasmic protein granules. Certain intrinsic features of granule components may also be relevant, such as potential multivalent interactions contributed by scaffolding and adaptor proteins (e.g. GRB2) and altered structural and biophysical properties of established cancer-derived mutants of RAS proteins (Hunter et al., 2015). There is clinical relevance of these future studies, as activating oncogenic mutations in RAS and other RAS/MAPK pathway components can co-occur with RTK gene rearrangements, and are also drivers of acquired resistance to kinase inhibitors in RTK fusion cancers (Kato et al., 2017; Pietrantonio et al., 2017; Rotow and Bivona, 2017). The interplay between cytoplasmic protein granule-based and canonical lipid membrane-based RTK/RAS/MAPK signaling will be important to understand for current and future therapeutic strategies that target oncogenic RTKs.

In summary, we report on the discovery of a cancer-specific membraneless subcellular structure formed by higher-order assembly of an RTK oncoprotein that is critical for oncogenic RTK/RAS signaling. Our results provide rationale for a new class of targeted therapeutics that aim to disrupt protein granule assembly and function. Treatment for oncogenic RTK/RAS/MAPK driven cancers is almost universally characterized by the development of drug resistance to targeted kinase inhibitors (Rotow and Bivona, 2017). Identifying critical factors that regulate the nucleation and degradation of RTK membraneless cytoplasmic protein granules, as well as defining roles for molecular chaperones and signaling proteins that promote multivalency-driven condensate formation, may provide opportunities for the development of a distinct class of targeted drugs to disrupt protein granules that drive cancer pathogenesis.

## Acknowledgments

The authors would like to acknowledge Amit Sabnis, Franziska Haderk and Zoji Bomya for experimental help and manuscript review, Mark Philips, Ignacio Rubio, and Richard Bayliss for generously providing plasmids and manuscript review, and Michael Rosen for scientific input and manuscript review.

## Author Contributions

A.T., J.G., D.S.N., B.H. and T.G.B. designed the study. A.T., J.G., D.S.N., H.L., A.H., H.A., S.P., A.R., X.S., B.Y., S.F. performed experiments, collected and analyzed data. A.T., J.G., D.S.N., B.H. and T.G.B wrote the manuscript. B.H. and T.G.B oversaw the study. All authors have approved the manuscript.

## Data and Materials Availability

All data is available in the main text or the supplementary materials.

## Supplementary Materials

### Supplementary Materials

STAR* Methods

Video S1 Description

Supplementary Figures 1 to 7

### STAR* Methods

#### Key Resource Table

| Reagent or Resource | Source | Identifier |
| --- | --- | --- |
| **Antibodies** |  |  |
| ALK (D5F3) | Cell Signaling Technology | Cat #3633 |
| p-Y1604-ALK | Cell Signaling Technology | Cat #3341 |
| ERK1/2 | Cell Signaling Technology | Cat #3493 |
| p-T202/Y204-ERK1/2 | Cell Signaling Technology | Cat #4370 |
| DCP1B (D2P9W) | Cell Signaling Technology | Cat #13233 |
| EEA1 (C45B10) | Cell Signaling Technology | Cat #3288 |
| EGF Receptor (D38B1) | Cell Signaling Technology | Cat #4267 |
| p-Y1068-EGF Receptor (D7A5) | Cell Signaling Technology | Cat #3777 |
| MEK1/2 | Cell Signaling Technology | Cat #9122 |
| p-S221-MEK1/2 (166F8) | Cell Signaling Technology | Cat #12338 |
| RET (C31B4) | Cell Signaling Technology | Cat #3223 |
| p-Y905-RET | Cell Signaling Technology | Cat #3221 |
| GFP/YFP (D5.1) | Cell Signaling Technology | Cat #2956 |
| HA (C29F4) | Cell Signaling Technology | Cat # 3724 |
| HRP-conjugated anti-mouse | Cell Signaling Technology | Cat #7076 |
| HRP-conjugated anti-rabbit | Cell Signaling Technology | Cat #7074 |
| Actin | Santa Cruz Biotechnologies | Cat # sc-1616 |
| HRAS | Santa Cruz Biotechnologies | Cat # sc-520 |
| NRAS | Santa Cruz Biotechnologies | Cat # sc-31 |
| KRAS | Santa Cruz Biotechnologies | Cat # sc-30 |
| Calnexin | Santa Cruz Biotechnologies | Cat # sc-23954 |
| GRB2 | Santa Cruz Biotechnologies | Cat # sc-8034 |
| Anti-Ras, clone 10 | EMD-Millipore | Cat #05-516 |
| Anti-FLAG M2 monoclonal | Sigma-Aldrich | Cat #F3165 |
| Alexa Fluor 488/594 | Abcam | Cat #ab150113,<br>Cat #ab150080 |
| **Chemicals, Peptides, and Recombinant Proteins** |  |  |
| Crizotinib | Selleck Chemicals | Cat #S1068 |
| Erlotinib | Selleck Chemicals | Cat #S7786 |
| Triton X-100 | Sigma-Aldrich | Cat #T8787 |
| RNase-A | Thermo Fisher Scientific | Cat #12091021 |
| HALT Protease inhibitor cocktail | Thermo Fisher Scientific | Cat #87785 |
| HALT Phosphatase inhibitor cocktail | Thermo Fisher Scientific | Cat #78420 |
| DMSO | Sigma-Aldrich | Cat #D2650 |
| 1,6-Hexanediol | Sigma-Aldrich | Cat #240117 |
| **Critical Commercial Assays** |  |  |
| RAS GST-RBD Activation Kit | Cytoskeleton | Cat #BK008 |
| MycoAlert Mycoplasma Detection Kit | Lonza | Cat #LT07-118 |
| Amersham ECL Prime Western Blotting Detection Reagent | GE Life Sciences | Cat #RPN2232 |
| Mirus Bio TransIT-LT1 | Fisher Scientific | Cat #MIR 2304 |
| M2 agarose flag beads | Sigma Aldrich | Cat #A2220 |
| CellTracker CM-DiI Membrane Dye | Thermo Fisher Scientific | Cat #C7001 |
| Anti-HA beads | Thermo Fisher Scientific | Cat #88836 |
| **Experimental Models: Cell Lines** |  |  |
| Beas2B | ATCC | CRL-9609 |
| H3122 | Generous gift from Christine Lovly | N/A |
| STE-1 | Generous gift from Christine Lovly | N/A |
| HCC827 | ATCC | Cat #CRL-2868 |
| HEK 293T cells | ATCC | Cat #CRL-11268 |
| **Oligonucleotides** |  |  |
| Sequences for sgRNA | This paper, see “cell line generation” | N/A |
| **Software and Algorithms** |  |  |
| Graphpad Prism 6 | Graphpad software | https://graphpad.com |
| ImageJ | Schneider et al., 2012 | https://imagej.nih.gov/ij |
| Micro-Manager software | Edelstein et al, 2010 | https://micro-manager.org |
| CellProfiler software | CellProfiler | http://cellprofiler.org |
| MatLab software | Mathworks | https://www.mathworks.com/products/matlab.html |

#### Lead Contact

Further information and requests for resources should be directed to and will be fulfilled by the Lead Contact, Trever Bivona.

#### Material Availability

Plasmids generated in this study are available by request from the Lead Contact.

#### Data and Code Availability

Data and codes generated in this study are available by request from the Lead Contact.

### Experimental Model and Subject Details

#### Cell Lines

This study utilized Beas2B, H3122, STE-1, HCC827, and 293T cells. All cell lines were maintained in humidified incubators with 5% CO2 at 37 °C. Beas2B and endogenously tagged derivatives, as well as the patient derived cancer cell lines H3122, STE-1, and HCC827 were cultured in RPMI-1640 medium supplemented with 10% FBS and penicillin-streptomycin at 100 µg/mL. 293T cells were cultured in DMEM-High Glucose supplemented with 10% FBS and 100 µg/mL of penicillin/streptomycin. All cell lines were tested for mycoplasma every 3 months using MycoAlert Mycoplasma Detection Kit (Lonza, Basel, Switzerland). All cells used were <20 passages from thaw.

### Method Details

#### Cell line generation

Generation of endogenously tagged mNeonGreen2_1-10/11_ cell lines was performed in the human bronchial epithelial cell line (Beas2B) and the patient-derived oncogenic ALK cell line (H3122) as previously described (Feng et al., 2017). Correct integration of mNeonGreen2_11_ was confirmed by genomic sequencing and by reduction in fluorescence upon gene knockdown. sgRNA spacer sequences used in this study are shown below.

GRB2: CTTAGACGTTCCGGTTCACG SOS1: ACAGAGGAACTCAGGAAGAA GAB1: GCGAAACCGTCCATCTTGCG KRAS: AATGACTGAATATAAACTTG HRAS: GATGACGGAATATAAGCTGG NRAS: AATGACTGAGTACAAACTGG

#### Biochemical fractionation

Cells were seeded in 10 cm dishes and harvested the following day by scraping into buffer A [10 mM Tris-HCl (pH 7.4), 1 mM EDTA, 250 mM Sucrose, supplemented with 1X HALT protease inhibitors (Thermo Fisher Scientific)]. Lysates were gently sonicated on minimum intensity and cleared by centrifugation. Lysate was then split equivalently to two ultracentrifugation tubes, and one tube was supplemented with 1% Triton X-100. Lysates were then subjected to ultracentrifugation at 100,000×g for 1 hour at 4 °C in an Optima MAX Ultracentrifuge (Beckman Coulter, Brea, CA). Supernatant and pelleted fractions were separated, resuspended with Laemmli sample buffer, boiled, and analyzed by SDS-PAGE. For RNase A experiments, lysates were incubated +/- RNase A at 1 µg/µL for 30 minutes at room temperature and then subjected to ultracentrifugation and processing as above.

#### Antibodies and immunoblotting

Antibodies against the following were obtained from Cell Signaling Technology (Danvers, MA, USA) and were used at a dilution of 1:1000: ALK (D5F3), p-Y1604-ALK (#3341), ERK1/2 (#3493), p-T202/Y204-ERK1/2 D13.14.4e (#4370), DCP1B (D2P9W), EEA1 (C45B10), EGF Receptor (D38B1), p-Y1068-EGF Receptor (D7A5), MEK1/2 (#9122), p-S221-MEK1/2 (#166F8), RET (C31B4), p-Y905-RET (#3221), GFP/YFP (D5.1) (#2956), HA (C29F4), horseradish peroxidase (HRP)-conjugated anti-mouse (#7076) and HRP-conjugated anti-rabbit (#7074). Antibodies to the following were obtained from Santa Cruz Biotechnologies (Santa Cruz, CA, USA): actin (I19, 1:1000 dilution), HRAS (C-20, 1:200 dilution), NRAS (F155, 1:200 dilution), KRAS (F234, 1:500 dilution), Calnexin Antibody (AF18), GRB2 (C7: 1:1000). Anti-Ras antibody, clone 10 (1:1000) was obtained from EMD Millipore (Hayward, CA) and anti-FLAG M2 monoclonal antibody was obtained from Sigma (USA).

For immunoblotting, cells were serum-starved (0% serum for 24 hours), then washed with ice-cold PBS and scraped in ice cold RIPA buffer [25 mM Tris⋅HCl (pH 7.6), 150 mM NaCl, 1% NP-40, 1% sodium deoxycholate, 0.1% SDS, supplemented with 1X HALT protease inhibitor cocktail and 1X HALT phosphatase inhibitor cocktail (Thermo Fisher Scientific)]. Lysates were clarified with sonication and centrifugation. Lysates were subject to SDS/PAGE followed by blotting with the indicated antibodies. Signal was detected using Amersham ECL Prime reagent (GE Healthcare Life Sciences, Chicago, IL, USA) and chemiluminescence on an ImageQuant LAS 4000 (GE Healthcare Life Science, Chicago, IL, USA).

#### Generation of stable cell lines expressing wild-type or cytosolic RAS

293T cells were infected with wild-type H/N/KRAS or respective cytosolic RAS mutants (KRAS C185S, HRAS C186S, NRAS C186S) then selected with puromycin to generate stable cell lines. RAS activation assays were performed 48 hours after transfection of empty vector, EML4-ALK, or oncogenic EGFR (EGFR L858R). H3122 and HCC827 cell lines were also infected with wild-type and C185S KRAS and selected with puromycin to generate stable cell lines. RAS activation assays were performed comparing 2 hour treatment with 100 nM crizotinib in H3122 cells or 100 nM erlotinib in HCC827 cells with mock DMSO treatment.

#### Compounds

Crizotinib and erlotinib were purchased from Selleck Chemicals (Houston, TX) and resuspended in DMSO.

#### RAS activation assays

The RAS GST-RBD activation kit was obtained from Cytoskeleton (Denver, CO, USA; #BK008). The protocol was according to the manufacturer’s instructions. Lysis buffer for RAS-GTP pulldowns was 50 mM Tris (pH 7.5), 10 mM MgCl_2_, 0.5 M NaCl, and 2% Igepal. 150 µg of lysate was incubated with 10µl RBD-beads overnight, followed by Western blotting. RAS-GTP levels were normalized to total RAS protein levels.

#### Live-cell microscopy

Cells were seeded in 35mm glass-bottom dishes (MatTek, Ashland, MA) or 8-well Nunc Lab-Tek 8 imaging chambers (Thermo Fisher Scientific) and then imaged using a Nikon Ti microscope with a CSU-W1 spinning disk confocal using a 100X/1.4 Plan Apo VC objective (Nikon Imaging Center, UCSF). Images were acquired on MicroManager software and analyzed using ImageJ software (Edelstein et al., 2014; Schneider et al., 2012).

#### Structured illumination microscopy

Beas2B cells were seeded into Nunc Lab-Tek 8-well imaging chambers and eYFP::EML4-ALK was transfected via Mirus TransIT-LT1 (Mirus Bio LLC, Madison, WI, USA). 24 hours later, both live cells and fixed cells were imaged with structured illumination microscopy on a DeltaVision OMX imaging system (GE Healthcare) in 3D-SIM mode. The procedure for cell fixation was 4% paraformaldehyde incubation for 5 minutes followed by three PBS washes. The 3D structures of granules were rendered with visualization software Chimera X.

#### Fluorescence recovery after photobleaching

For photobleaching experiments, Beas2B cells were seeded into Nunc Lab-Tek 8-well imaging chambers and transfected with mTagBFP2::EML4-ALK via Mirus TransIT-LT1. A 473 nm laser (Rapp Optoelectronic) at excitation intensity of 30 mW was used to photobleach regions of interest (ROIs) corresponding to individual granules in the sample. The fluorescence intensity was monitored before and after photobleaching with time interval of 3 seconds. Further intensity analyses were done in MatLab with custom-written code.

#### Monitoring granules during hexanediol treatment

Beas2B cells expressing mTagBFP2::EML4-ALK and GFP::DCP1B were seeded into Nunc Lab-Tek 8-well imaging chambers. A custom-made sample holder ensured the imaging chamber fits securely on the microscope stage without position shift. The cells were first imaged in regular RPMI cell culture media and then the media was replaced by RPMI media containing 5% hexanediol (Sigma). The same field of view was monitored at defined time points after addition of hexanediol.

#### Immunoprecipitation

For immunoprecipitation assays, HEK 293T cells were transfected with FLAG-tagged versions of EML4-ALK and respective mutants or HA- and YFP-tagged versions of EML4-ALK variants 1, 3, and 5. 48 hours post-transfection (after serum starvation in 0% serum for 24 hours), the cells were resuspended in lysis buffer (0.5% NP-40, 150 mM NaCl, 50 mM TrisCl, pH 7.5) containing protease and phosphatase inhibitor cocktails (Sigma). Lysates were syringed and centrifuged to clarify, then whole-cell extracts were either incubated overnight at 4°C with M2 agarose-FLAG beads (Sigma) or for one hour with anti-HA beads (Thermo Fischer Scientific). The immunocomplexes were washed three times with wash buffer (50 mM Tris (pH 7.4, 150 mM NaCl) and FLAG or HA beads were boiled and loaded for SDS-PAGE.

#### Immunofluorescence

H3122 or Beas2B cells expressing EML4-ALK were seeded in 4-well Lab Tek II Chamber Slides (Thermo Fisher Scientific). The following day, cells were fixed for 15 minutes with 4% paraformaldehyde, washed, and incubated in blocking buffer for 1 hour (1X PBS with 1% BSA and 0.3% Triton-X100). Blocking buffer was aspirated and cells were incubated with primary antibody (either ALK DF53 1:1000 from Cell Signaling Technology or FLAG-M2 1:1000 from Sigma) overnight in the dark at 4 °C. The following day, cells were washed, incubated with fluorophore-conjugated secondary antibodies (Alexa Fluor 488/594 from Abcam, 1:2000) for 1 hour at room temperature in the dark, washed, and then mounted using ProLong Gold Antifade reagent with DAPI (Cell Signaling Technology). Slides were analyzed using a Nikon Ti microscope with a CSU-W1 spinning disk confocal using a 100 × 1.4 NA Plan Apo VC objective (Nikon Imaging Center, UCSF). Images were acquired on MicroManager software and analyzed using ImageJ software.

#### Plasmids and construct generation

EML4-ALK cDNA and respective mutants were cloned into pBabe-puro with a N-terminal FLAG tag and mTagBFP2-C1. HA and YFP-tagged EML4-ALK variants 1, 3, and 5 were kind gifts from Dr. Richard Bayliss. EGFRL858R was cloned into pBabe-puro and mTagBFP2-N1. iEGFR was constructed from full length EGFR (NM_005228.5) by PCR amplification using the following primers: ATGcgaaggcgccacatcgttcgg and gtgaatttattggagca. HOtag sequences for forced clustering were provided by Dr. Xiaokun Shu (Zhang et al., 2018). The RAS-GTP reporters (tandem GFP-RBD and GFP-RBD mutant) were a kind gift from Dr. Ignacio Rubio. cDNA sequence for GRB2 was cloned into the mEGFP-C1 vector. All mutants were generated through a combination of QuikChange site-directed mutagenesis (Agilent) and gene synthesis (Genewiz). The mutants contain the following modifications:

EML4-ALK full-length sequence GenBank: AB274722.1, using cDNA from bases 271 – 3450, mutations/deletions based on this sequence numbering. EML4-ALK kinase-deficient K589M: A2036T, EML4-ALK ΔHELP: deletion of bases 928-1158, EML4-ALK ΔTD: deletion of bases 310-459. CCDC6-RET full-length sequence GenBank: KU254649.1, cDNA from bases 1-1512, mutations/deletions based on this sequence numbering. CCDC6-RET kinase-deficient K147M: A440T, CCDC6-RET ΔCC: deletion of bases 160-303.

#### DNA transfections

293T and Beas2B cells were transiently transfected using Mirus TransIt-LT1 transfection reagent according to manufacturer’s protocol.

#### Viral transduction

cDNAs for EML4-ALK, KRAS, HRAS, NRAS and respective cytosolic mutants were cloned into pLV-EF1a-IRES-blast (or hygromycin/puromycin selectable equivalent vectors). HEK 293T viral packaging cells were plated in 10 cm dishes the day prior to transfection. They were transfected with lentiviral or retroviral expression constructs and the appropriate packaging plasmids using Mirus TransIt-LT1 transfection reagent. Viral supernatants were collected 48-72 hours post-transfection and used to transduce cell lines in the presence of 1× Polybrene for 24 hours.

### Quantification and Statistical Analysis

#### Quantification of colocalization between EML4-ALK granules and endogenous signaling proteins

Customized MatLab code was written to correct for uneven illumination pattern in the optical path and cell autofluorescence background. Granules were identified with Cellprofiler feature-finding module using Otsu thresholding method and size constraint from 0.4 to 2 µm in diameter. The pair-wise centroid distance between features in BFP and GFP channels, corresponding to EML4-ALK granules and signaling proteins respectively, were calculated to identify colocalization events. Typically, ∼20 images containing 30+ cells and ∼300 granules for each signaling protein were analyzed in an automated batch-processing format. All colocalization events were confirmed manually by overlaying identified features with raw images.

#### Quantification of enrichment level of signaling proteins at EML4-ALK granules

Customized MatLab code was written to identify pixels corresponding to EML4-ALK granules in the BFP channel in a given image. The intensity in the GFP channel at the positions of these pixels, corresponding to the enriched signaling proteins, was averaged for individual granules. The ratio of the average pixel intensity at the granule to the pixel intensity averaged over the whole cell area is the fold enrichment of the signaling protein at each granule.

#### Quantification of fraction of granule containing cells for EML4-ALK and CCDC6-RET mutants

Beas2B cells were seeded into Nunc Lab-Tek 8-well imaging chambers and plasmids encoding wild-type and mutant forms of mTagBFP2::EML4-ALK (i.e. ΔTD, ΔHELP, K589M, see text for details), mTagBFP2::CCDC6-RET (i.e. ΔCC, K147M), as well as YFP::EML4-ALK variants 1, 3, and 5 were transfected using Mirus TransIT-LT1. 24 hours later, an initial position in each well was randomly picked as the center of an area of 650 µm x 650 µm and imaged with automated scanning and tiling done through MicroManager with a 100x oil objective (N.A. = 1.40). The process was repeated three times and all the cells were scored manually to determine if they contained cytoplasmic granules.

### Statistical analysis

*P* values were determined with Student’s *t*-tests or one-way ANOVA between comparator groups using GraphPad software.

### Supplementary Data

**Video S1:** Live-cell confocal microscopy video of Beas2B cells expressing tagBFP2::EML4- ALK showing EML4-ALK granules in direct contact without any evidence of fusion or fission events.

### Supplementary Figures 1-7 on pages 34-48

**Supplementary Figure 1:**
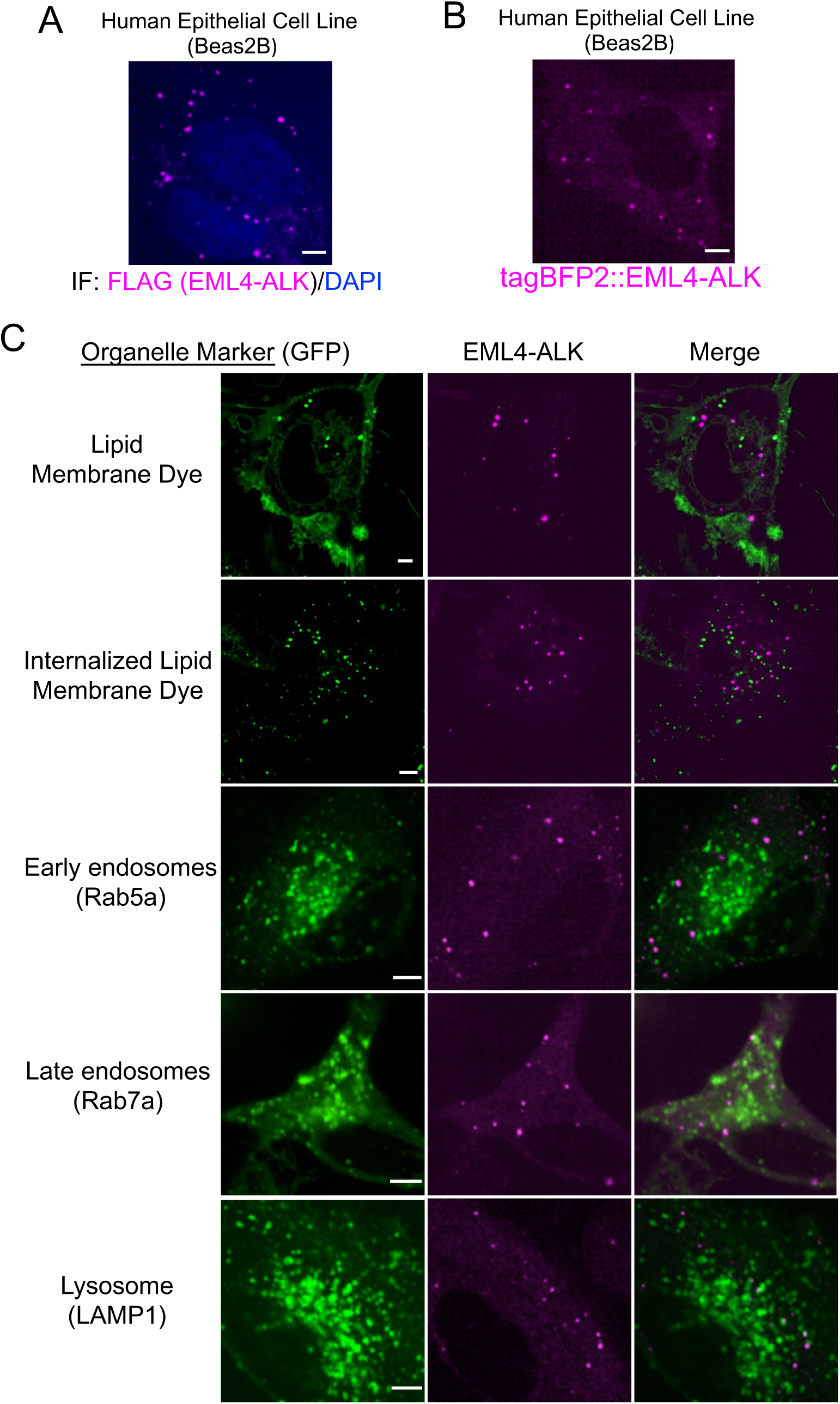
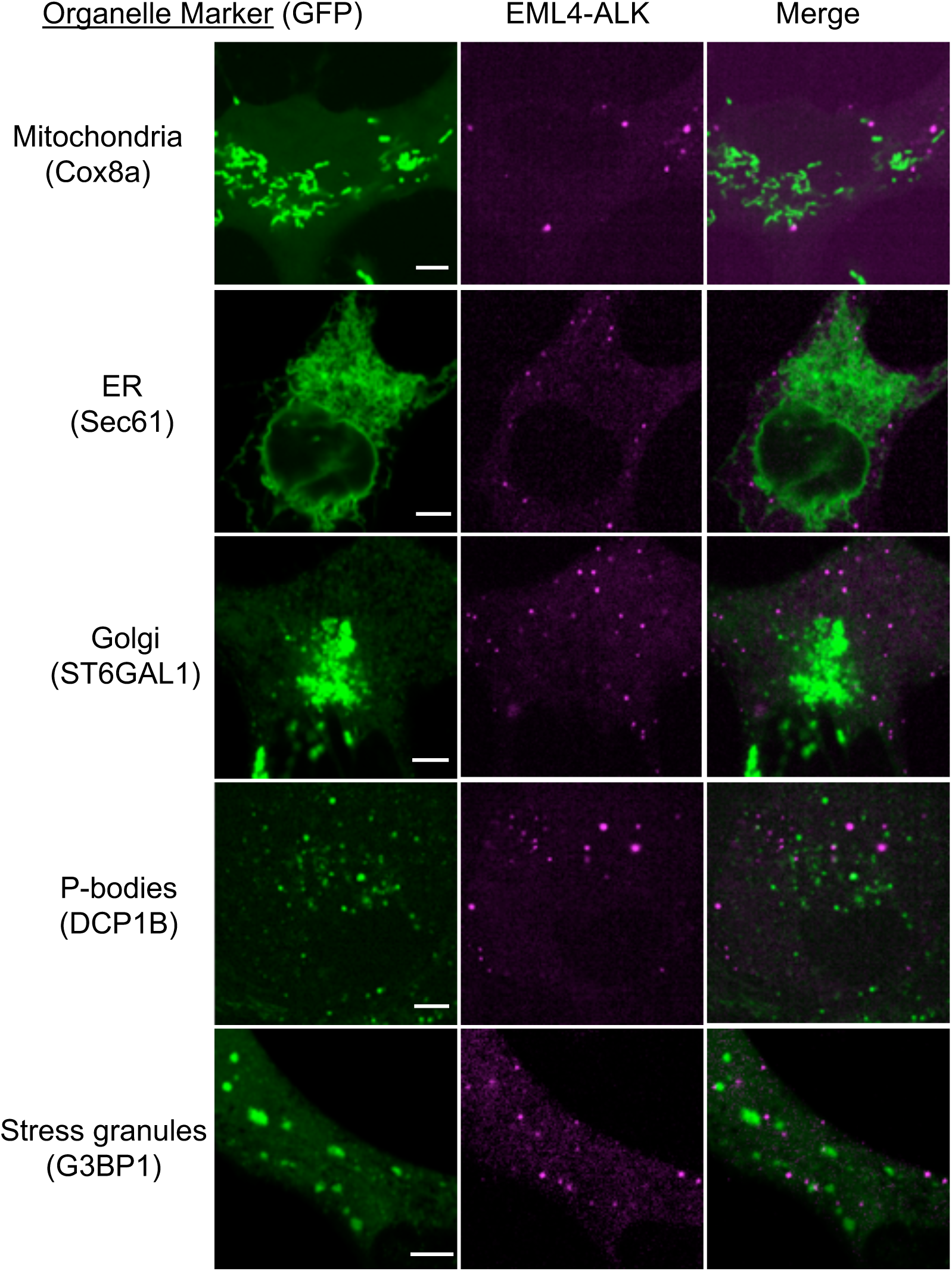
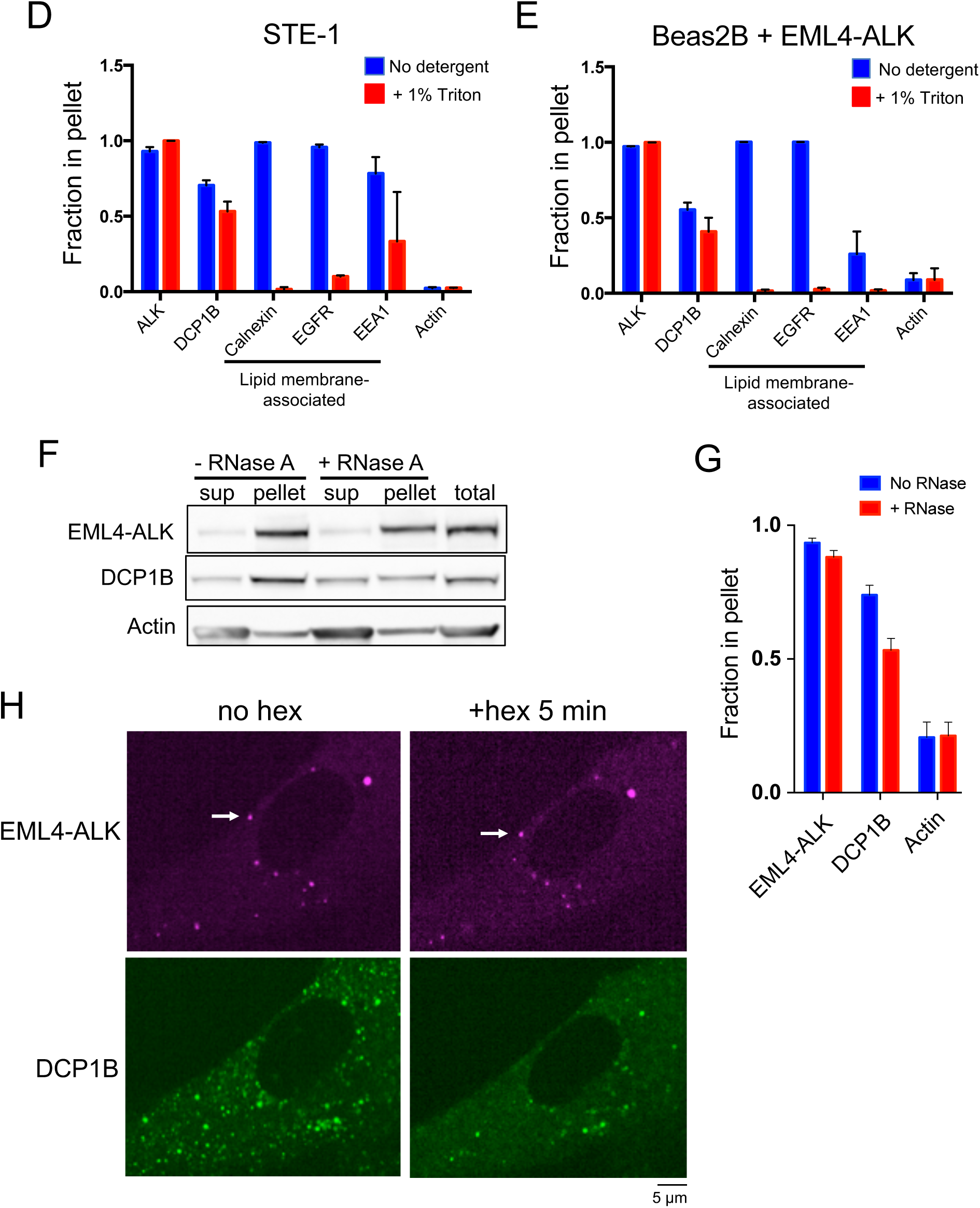
Cell biological and biophysical properties of EML4-ALK membraneless cytoplasmic protein granules. **(A)** Anti-FLAG immunofluorescence of FLAG-tagged EML4-ALK expressed in human epithelial cell line Beas2B. DAPI serves as a nuclear stain. Image is representative of at least 75 analyzed cells in total over 3 independent experiments. Scale bar = 5 µM. **(B)** Live-cell confocal imaging of human epithelial cell line Beas2B upon expression of mTagBFP2::EML4-ALK. Image is representative of 200 analyzed cells in total over 5 independent experiments. Scale bar = 5 µM. **(C)** Live-cell confocal imaging of human epithelial cell line Beas2B upon expression of mTagBFP2::EML4-ALK and mEGFP-tagged organelle markers as listed. Membrane dye experiments were conducted using live cells incubated with CellTracker™ CM-DiI Dye (Invitrogen). Each panel is a representative image of at least 20 analyzed cells per condition in 3 independent experiments. Scale bar = 5 µM. **(D, E)** Subcellular fractionation by ultracentrifugation +/– detergent (1% Triton X-100) to disrupt lipid membranes in STE-1 (D), an EML4-ALK expressing cancer cell line, and Beas2B cells expressing EML4-ALK (E). In both cell lines, EML4-ALK and DCP1B are statistically distinct from the lipid membrane-associated proteins, which shift from the insoluble fraction (pellet) to the soluble fraction with detergent (p < 0.05 for all comparisons by one-way ANOVA, except for DCP1B vs. EEA1 in panel D and E and ALK vs. EEA1 in panel E). Bar graphs reflect quantification of Western blotting results for 3 independent replicates. Fraction in pellet calculated as ratio of the insoluble fraction to total (insoluble plus supernatant fractions) as assessed by Western blotting. Error bars represent ± SEM. **(F, G)** Subcellular fractionation by ultracentrifugation +/- RNase A treatment for 30 minutes in EML4-ALK expressing cancer cell line H3122. P-body protein DCP1B partially shifts from the insoluble (pellet) fraction to the supernatant (sup) upon RNase A treatment, in contrast to EML4- ALK. Western blotting images are representative of at least 5 independent experiments. Fraction in pellet (G) calculated as ratio of the insoluble fraction to total (insoluble plus supernatant fractions) as assessed by Western blotting (F). DCP1B demonstrates a significant RNase-dependent reduction in the insoluble fraction (p < 0.05 by paired t-test) compared to EML4-ALK. **(H)** Live-cell imaging of human epithelial cell line Beas2B co-expressing mTagBFP2::EML4- ALK and eGFP::DCP1B treated with 5% hexanediol (hex) and imaged at respective time points. Images are representative of at least 5 analyzed cells in 3 independent experiments. White arrows indicate a representative EML4-ALK cytoplasmic protein granule (multiple non-highlighted granules are also pictured in all panels). Quantification of granule persistence shown in Figure 1D.

**Supplementary Figure 2:**
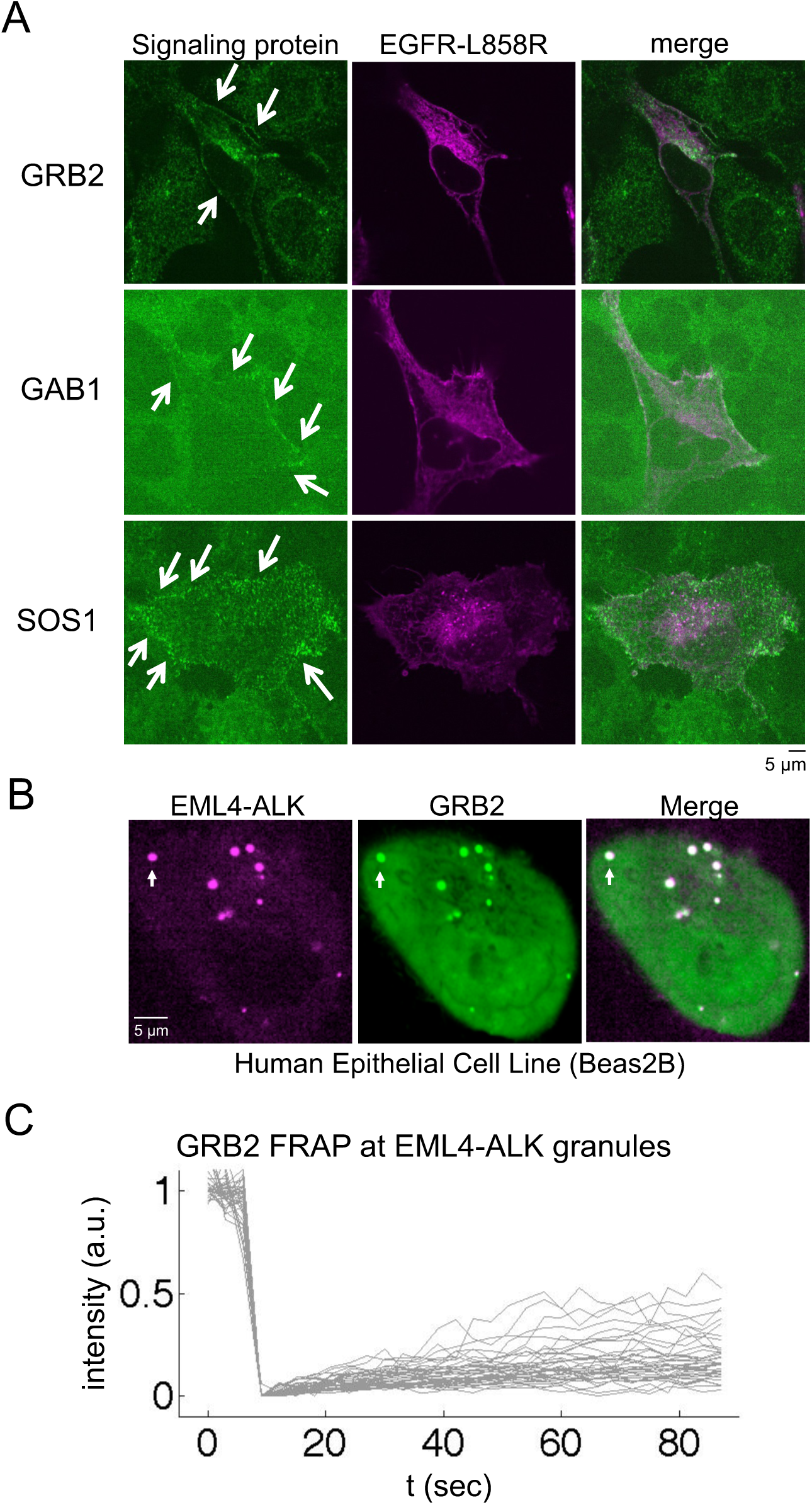
RAS adaptor protein GRB2 is recruited to EML4-ALK cytoplasmic protein granules. **(A)** Live-cell confocal imaging of mTagBFP2::EGFR L858R (oncogenic EGFR) expressed in human epithelial cell lines (Beas2B) with endogenous mNeonGreen2-tagging of GRB2, GAB1, and SOS1. Arrows denote plasma membrane enrichment of GRB2, GAB1, and SOS1. Representative images from at least 30 cells analyzed per condition in 3 independent experiments. **(B)** Live-cell confocal imaging of mTagBFP2::EML4-ALK and mEGFP::GRB2 upon dual expression in Beas2B cells. White arrows indicate a representative EML4-ALK cytoplasmic protein granule with local enrichment of GRB2 (multiple non-highlighted granules also show colocalization between EML4-ALK and GRB2). Images are representative of 100 analyzed cells in total over 3 independent experiments. **(C)** FRAP experiments performed in human epithelial cells (Beas2B) with endogenous mNeonGreen2-tagging of GRB2, upon expression of mTagBFP2::EML4-ALK. Graph displays individual recovery of fluorescence intensity after photobleaching of GRB2 enriched at EML4- ALK granules, t denotes time in seconds, intensity in arbitrary units (a.u.) normalized to 1. N = 30 cells.

**Supplementary Figure 3:**
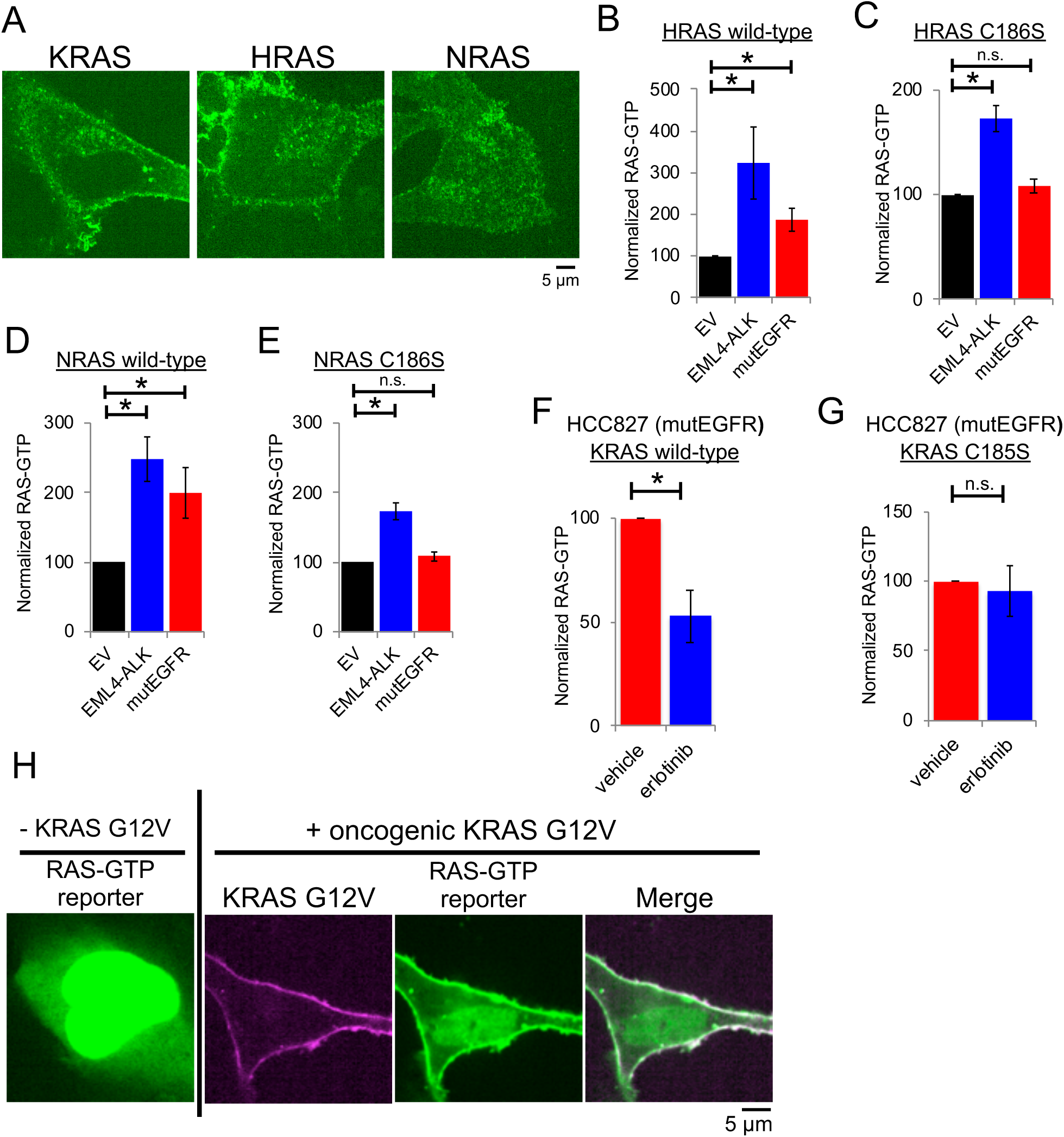
EML4-ALK-specific activation of cytosolic RAS. **(A)** Live-cell confocal imaging of human epithelial cell lines (Beas2B) with endogenous mNeonGreen2-tagging of KRAS, HRAS, and NRAS. Images are representative of at least 20 analyzed cells per condition in 3 independent experiments. **(B-E)** HRAS/NRAS wild-type or cytosolic mutants (HRAS C186S, NRAS C186S) were stably expressed in 293T cells and then transfected with either empty vector (EV), EML4-ALK, or oncogenic EGFR-L858R (mutEGFR). RAS-GTP levels were normalized to the relevant total RAS protein level (H/NRAS wild-type or C186S). N = 4. Error bars represent ± SEM, * denotes p value < 0.05, n.s. denotes non-significant comparison, one-way ANOVA. **(F, G)** Patient-derived oncogenic EGFR expressing cell line (HCC827) with stable expression of either wild-type KRAS or cytosolic mutant (KRAS C185S). RAS-GTP levels determined +/– two hours of 100 nM erlotinib treatment and normalized to the relevant total RAS protein level (KRAS wild-type or C185S). N = 4. Error bars represent ± SEM, * denotes p value < 0.05, n.s. denotes non-significant comparison, paired *t-*test. **(H)** Live-cell confocal imaging of human epithelial cell line Beas2B with GFP-labelled RAS-GTP reporter (tandem GFP-RBD). Left column demonstrates baseline RAS-GTP reporter localization to cytosol and nucleoplasm, right three panels show plasma membrane re- localization of RAS-GTP reporter upon expression of mTagBFP2::KRAS G12V (oncogenic KRAS). Images are representative of 15-25 cells analyzed per condition in 3 independent experiments.

**Supplementary Figure 4:**
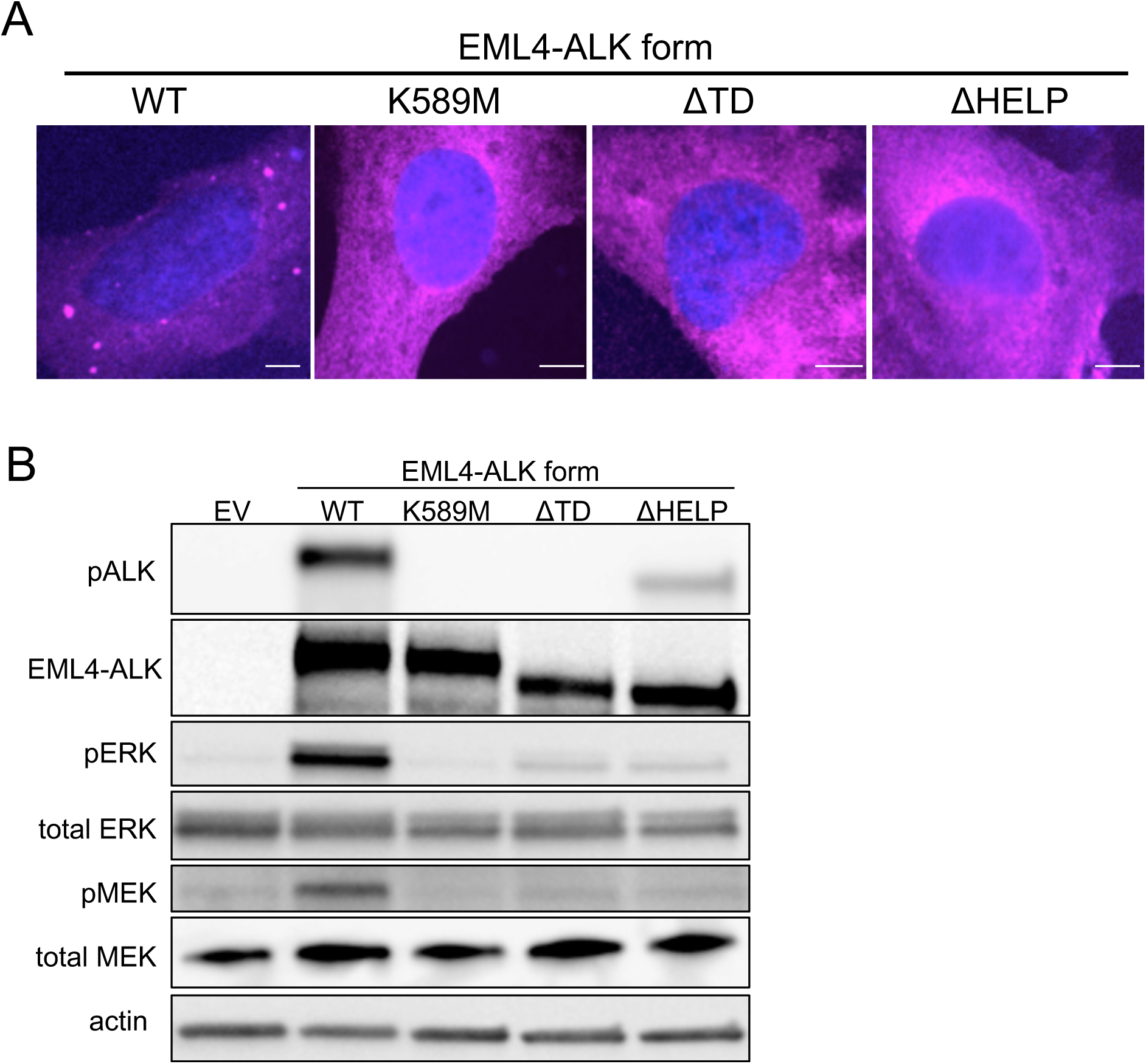
Non-granule-forming mutants of EML4-ALK disrupt RAS/MAPK signaling. **(A)** Anti-FLAG immunofluorescence of human epithelial cell line Beas2B expressing either FLAG-tagged EML4-ALK wild-type (WT) or EML4-ALK kinase-deficient (K589M), ΔTD or ΔHELP mutants. EML4-ALK (FLAG) staining in pink, DAPI in blue. Images are representative of at least 35 cells analyzed per condition in 2 independent replicates. Scale bar = 5 µM. **(B)** Western blotting upon expression of wild-type EML4-ALK or EML4-ALK kinase-deficient (K589M), ΔTD or ΔHELP mutants in 293T cells. Images are representative of at least 5 independent experiments and pERK levels are quantified in Main Figure 4F.

**Supplementary Figure 5:**
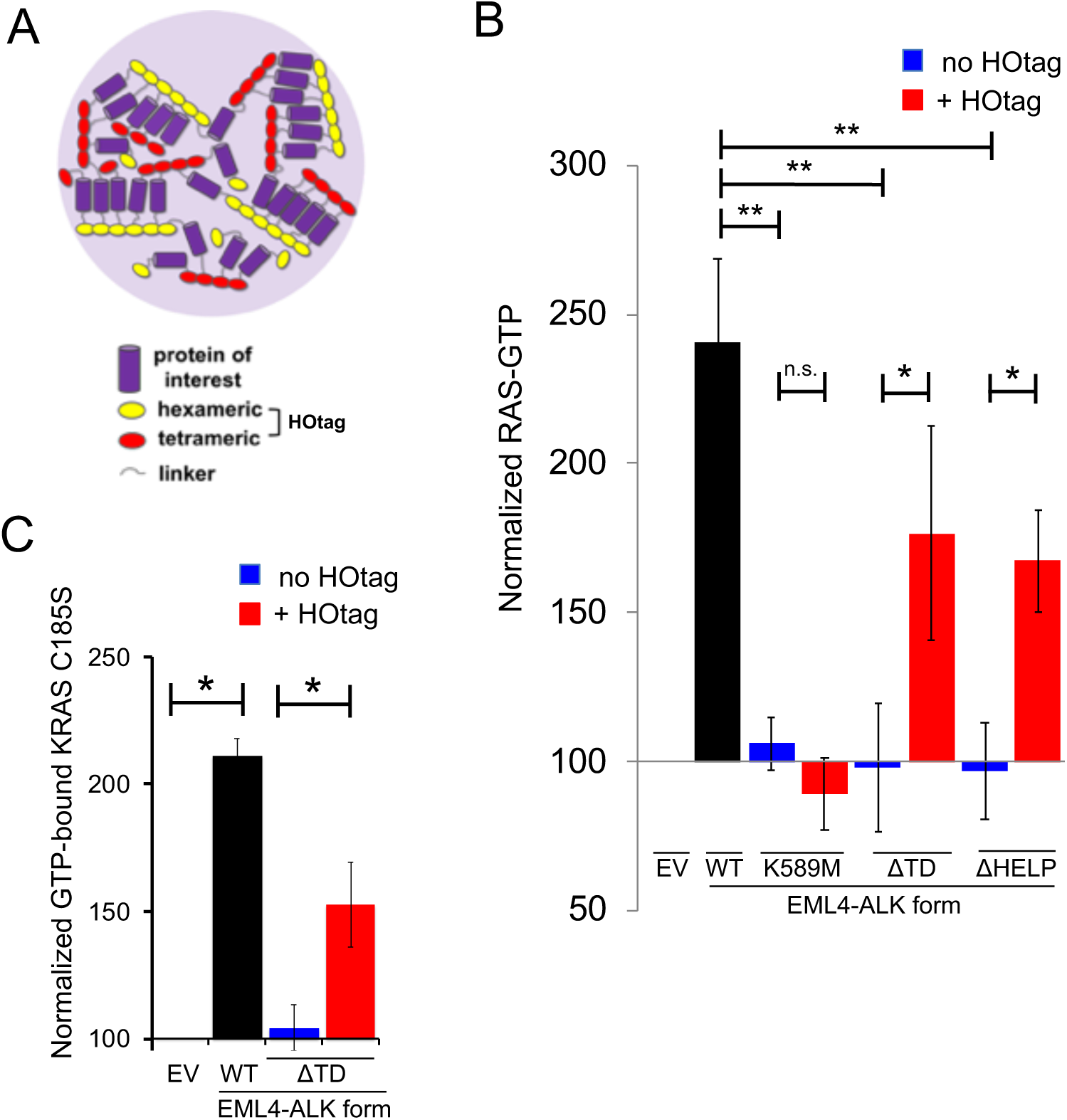
Forced granule formation of cytosolic EML4-ALK mutants rescues RAS-GTP levels. **(A)** Forced clustering of proteins achieved by N-terminal hexameric and C-terminal tetrameric tags that form higher-order clustered assemblies upon expression in cells. **(B)** Quantification of endogenous RAS-GTP levels performed in 293T cells expressing an empty vector (EV), EML4-ALK wild-type (WT), or the diffusely cytosolic EML4-ALK mutants (kinase deficient K589M, ΔTD, or ΔHELP) +/– forced clustering (HOtag). EML4-ALK K589M, ΔTD and ΔHELP mutants (blue bars) display significantly reduced RAS-GTP levels compared to wild-type EML4-ALK (black bar), ** denotes p < 0.01 by one-way ANOVA. Forced clustering (HOtag, red bars) of EML4-ALK ΔTD and ΔHELP mutants, but not EML4-ALK K589M, significantly increases RAS-GTP levels compared to the respective non-clustered EML4-ALK mutants (blue bars), * denotes p < 0.05, n.s. denotes non-significant comparison, paired *t*-test. RAS-GTP levels normalized to total RAS protein levels. N = 4. Error bars represent ± SEM. **(C)** Stable expression of cytosolic KRAS mutant (KRAS C185S) in 293T cells, followed by transfection of empty vector (EV), EML4-ALK wild-type (WT) or diffusely cytosolic EML4- ALK ΔTD mutant +/– forced clustering (HOtag). Levels of GTP-bound KRAS C185S were normalized to total KRAS C185S protein levels. N = 4. Error bars represent ± SEM, * denotes p < 0.05 by paired *t*-test.

**Supplementary Figure 6:**
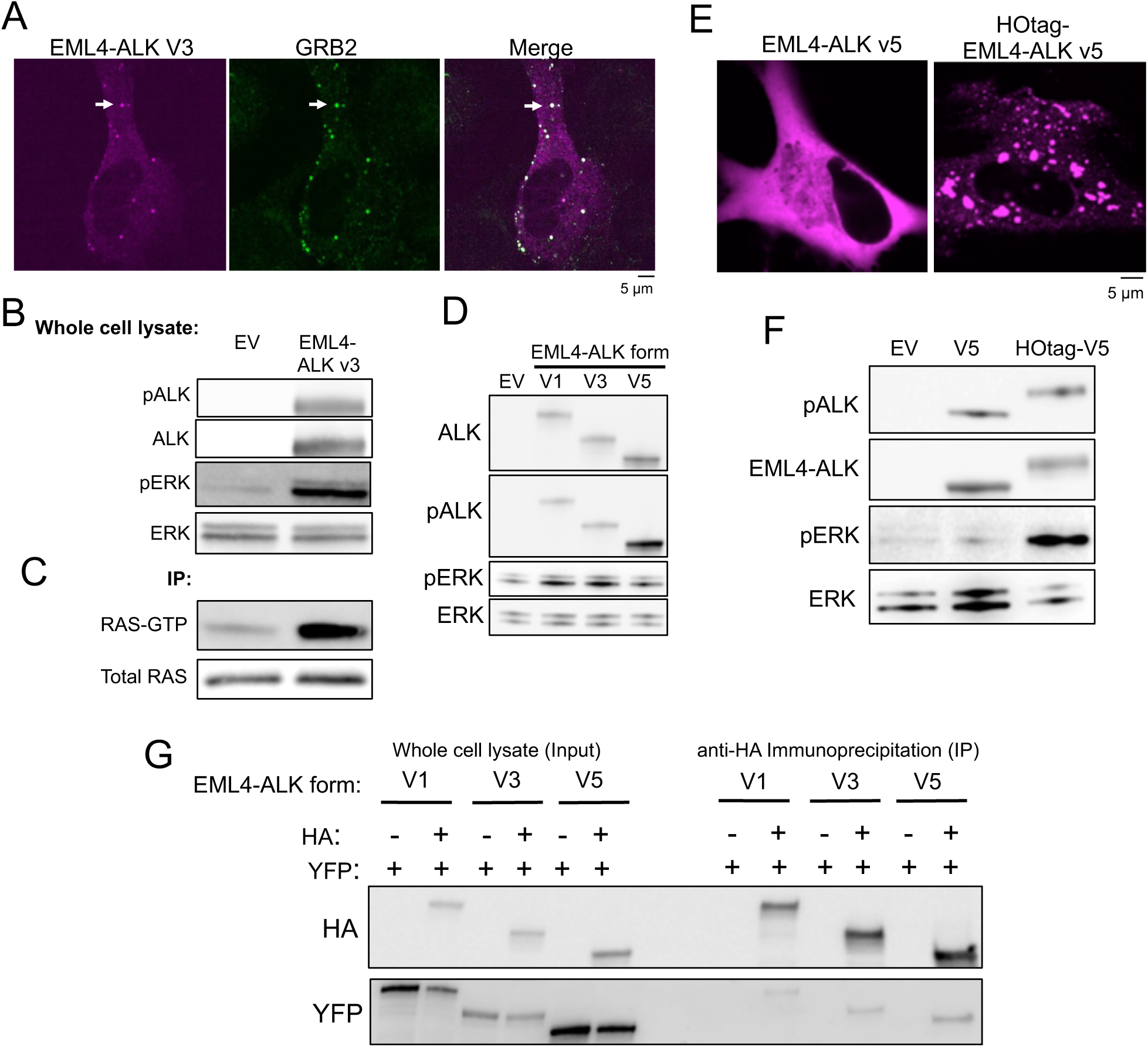
Higher-order protein assembly of RTKs increases RAS/MAPK signaling. **(A)** Live-cell confocal imaging of mTagBFP2::EML4-ALK variant 3 expressed in human epithelial cell line Beas2B with endogenous mNG2-tagging of GRB2. White arrows indicate a representative EML4-ALK variant 3 cytoplasmic protein granule with local enrichment of GRB2 (multiple non-highlighted granules also show colocalization between EML4-ALK variant 3 and GRB2). Images are representative of at least 25 analyzed cells in 3 independent experiments. **(B, C)** Western blot analysis upon expression of empty vector (EV) or EML4-ALK variant 3 in 293T cells to assess levels of EML4-ALK activation (phosphorylation), ERK activation, and in immunoprecipitation (IP) panel (C), levels of RAS activation (GTP-bound RAS). Representative images from at least 4 independent experiments. **(D)** Western blot analysis upon expression of empty vector (EV) or EML4-ALK variants 1, 3, or 5 in 293T cells to assess levels of ERK activation. Images are representative of at least 4 independent experiments and pERK levels are quantified in Main Figure 6D. **(E)** Live-cell confocal imaging of YFP::EML4-ALK variant 5 +/- forced clustering (HOtag) in human epithelial cell line Beas2B. Images are representative of at least 20 analyzed cells in 3 independent experiments. **(F)** Western blot analysis upon expression of empty vector (EV) or EML4-ALK variant 5 +/- HOtag in 293T cells. Images are representative of at least 4 independent experiments. **(G)** YFP- and HA-tagged versions of EML4-ALK variants 1, 3, and 5 were expressed in 293T cells as indicated. Western blot analysis of input and immunoprecipitation (IP) with anti-HA beads. Images are representative of 3 independent experiments.

**Supplementary Figure 7:**
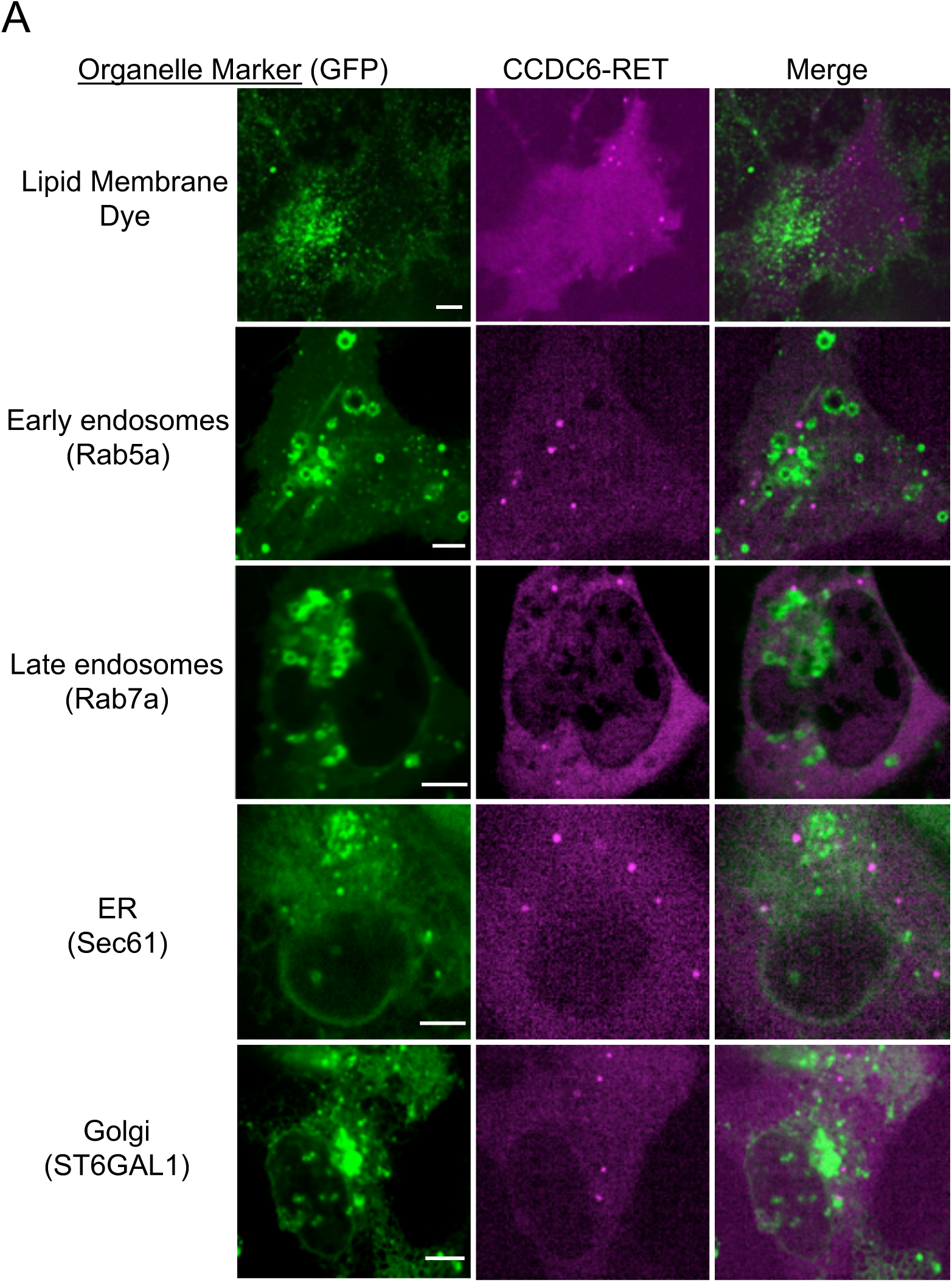
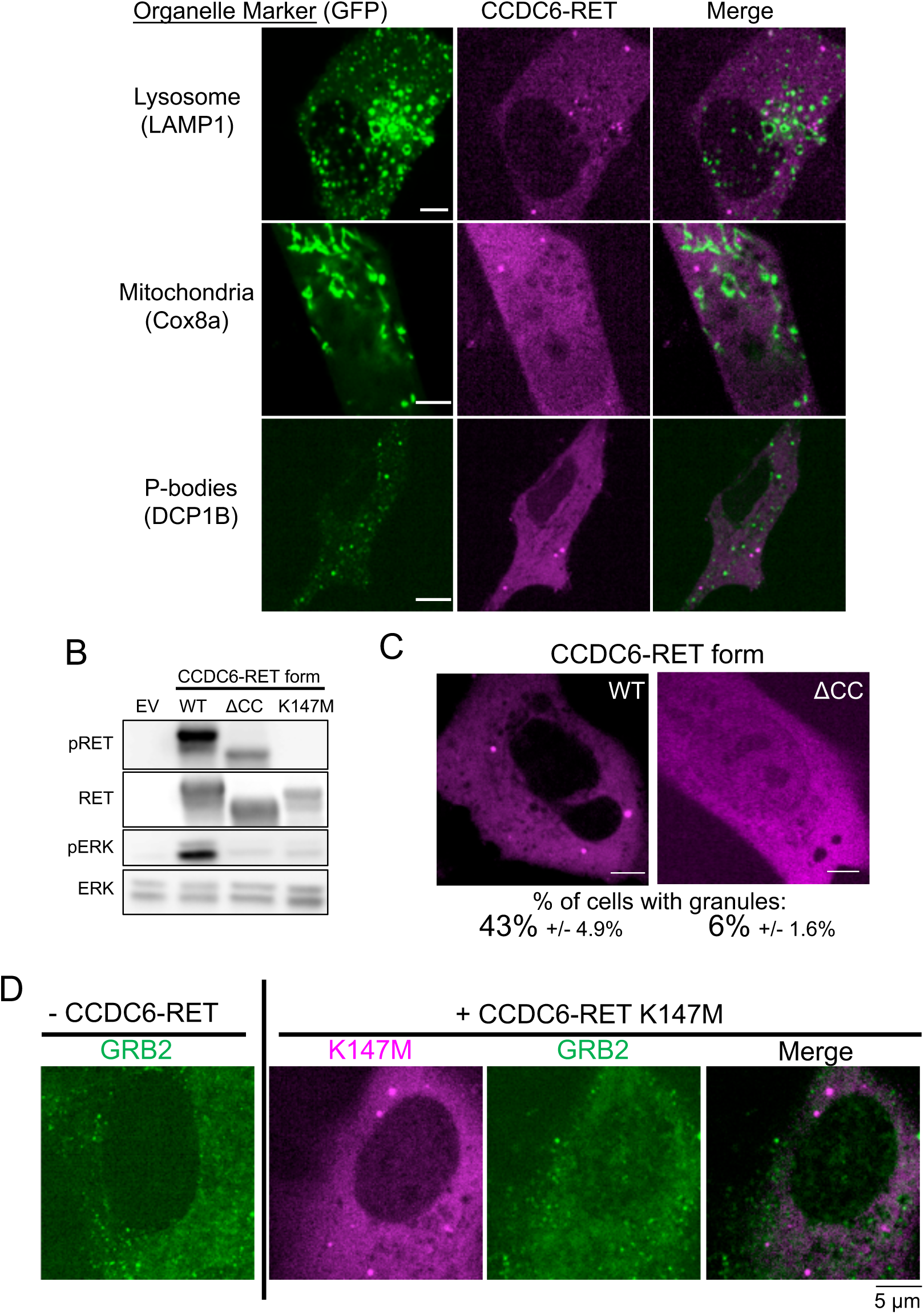
CCDC6-RET forms membraneless cytoplasmic protein granules that are critical for RAS/MAPK pathway activation. **(A)** Live-cell confocal imaging of human epithelial cell line Beas2B upon expression of mTagBFP2::CCDC6-RET and mEGFP-tagged organelle markers as listed. Membrane dye experiments were conducted using live cells incubated with CellTracker™ CM-DiI Dye (Invitrogen) according to manufacturer’s recommended protocol. Each panel is a representative image of at least 20 analyzed cells per condition with at least 3 independent replicates. **(B)** Western blotting upon expression of an empty vector (EV), CCDC6-RET wild-type (WT) or CCDC6-RET mutants (coiled-coiled domain deletion mutant, ΔCC, and kinase-deficient mutant K147M) in 293T cells. Western blot images are representative of 4 independent experiments and pERK levels are quantified in Main Figure 7E. **(C)** Live-cell confocal imaging of human epithelial cell line Beas2B expressing mTagBFP2- labelled CCDC6-RET wild-type (WT) or the coiled-coiled domain deletion mutant (ΔCC). Quantification of cells with 3 or more granules shown as a fraction ± SEM, based on 3 independent experiments of at least 25 cells analyzed per condition. Scale bar = 5 µM. **(D)** Live-cell confocal imaging of mTagBFP2::CCDC6-RET K147M (kinase-deficient mutant) expressed in the human epithelial cell line Beas2B with endogenous mNG2-tagging of GRB2. Images are representative of 60 analyzed cells in total over 3 independent experiments with no observed local enrichment of GRB2 at CCDC6-RET K147M granules.

